# Molecular basis of arthropod appendage diversity

**DOI:** 10.1101/2025.01.27.634880

**Authors:** Heather S. Bruce, Nipam H. Patel

**Affiliations:** University of British Columbia; Marine Biological Laboratory and University of Chicago

## Abstract

Arthropods have an incredible diversity of limbs that are modified for walking, chewing, cleaning, mating, grasping, sensing, and more. Understanding the relationships and evolutionary histories of different limbs is a central task, but their sheer diversity makes this a daunting if not impossible task using morphology alone. Here, the in situ expression patterns and CRISPR-Cas9 phenotypes for the five best-studied leg-patterning genes – *Distal-less*, *Sp6-9*, *dachshund*, *extradenticle*, and *homothorax* – are described for all limbs of the crustacean *Parhyale*.

Crustaceans are well-suited for this task because their limbs are more diverse than those of other arthropods, and each individual possesses a wide range of limb types that are relevant to many other arthropods, living and extinct. These results will a) provide a template for understanding the genetic basis of limb construction in arthropods more generally based on the strong phenotypes that can be obtained with CRISPR-Cas9, and b) contribute to our understanding of the evolution and affinities of highly modified legs like mouthparts and genitalia using molecular methods to complement previous morphological and embryological approaches.

## MAIN TEXT

Arthropods ancestrally possess 8 leg segments (podomeres)(G. A. Boxshall, 2004; Bruce, 2022; Bruce & Patel, 2020; Ewing, 1928; Shultz, 1989; Yang et al., 2018), but this condition has been modified in different arthropod lineages and in different arthropod limb types. Different arthropod lineages have incorporated one or more proximal leg segments into the body wall (Bruce, 2022; Bruce & Patel, 2020; Matsuda, 1963; Snodgrass, 1927; Tiegs, 1940) leading to a reduction in the number of freely moveable leg segments (6 leg segments in insects, 7 or 8 in crustaceans, 6 or 7 in myriapods, and 7 or 8 in chelicerates). Some arthropods have truncated the distal leg (Bruce & Patel, 2023). Furthermore, the highly modified morphologies of some limbs like the mouthparts, antennae, and genitalia make it difficult to understand how these correspond to the 8 ancestral leg segments (Boudinot, 2018; Coulcher & Telford, 2013; Klass & Matushkina, 2012; Popadić et al., 1998; Scholtz et al., 1998; Yang et al., 2018).

Given that all arthropod legs appear to follow the same molecular patterning plan (Bruce, 2022; Bruce & Patel, 2020, 2022, 2023) investigating the function of leg patterning genes in a representative arthropod will provide more general insight into the patterning mechanisms for other arthropod legs. Crustaceans are an ideal representative for this task because their limbs are more diverse than those of other arthropods and retain features that have been variably lost in other lineages. For example, the genetically-tractable amphipod crustacean, *Parhyale hawaiensis*, possesses a wide range of limb types that are relevant to many other arthropods, living and extinct: antennae, mandibles, maxillae, maxilliped, claws, walking legs, jumping legs, biramous swimmerets, and biramous anchor legs (uropods), as well as a labrum and paragnaths, and a variety of exites (lateral leg lobes such as gills or plates) and endites (medial leg lobes such as those on the mouthparts). The eight ancestral leg segments in crustaceans correspond to the *dac*tyl (1), propodus (2), carpus (3), merus (4), ischium (5), basis (6), coxa (7), precoxa (8).

Previous work on leg-patterning in *Parhyale* used CRISPR-Cas9 gene editing to knock out the five best-studied leg-patterning genes in arthropods: *Distal-less* (*Dll*), *Sp6-9*, *dachshund* (*dac*), *extradenticle* (*exd*), and *homothorax* (*hth*) and reported phenotypes for the third thoracic limb (T3) of *Parhyale*, a claw (cheliped) (Bruce, 2022; Bruce & Patel, 2020). Here, the phenotypes for all five leg-patterning genes in all limb types of *Parhyale* is reported for whole hatchlings using confocal microscopy, as well as dissected appendages using brightfield and darkfield microscopy.

### WT *PARHYALE* BODY PLAN

The *Parhyale* body plan is composed of a 6- or 7-segmented head, 8-segmented thorax, and 6-segmented abdomen with telson (Figs 1, 2) (Browne et al., 2005). Each leg has a unique suite of characters that readily distinguishes it from the others. The head is composed of a labrum of unknown affinity (Lb, Fig. 1; (Ortega-Hernández & Budd, 2016; Scholtz & Edgecombe, 2006)), an ocular segment (and pre-ocular region which may or may not be a “true segment” (Marlow et al., 2014; Posnien et al., 2023; Schmidt-Ott & Technau, 1992)), first antenna (An1), second antenna (An2), mandible (Md) with paragnath (pg), first maxilla (Mx1), and second maxilla (Mx2). All head appendages are tightly compacted together and highly modified, making it difficult to assign leg segment (podomere) homologies based on morphology alone. The labrum is a fused pair of buds that functions as an “upper lip”; it arises in the embryo anterior to An1 and then migrates ventrally to nestle between An2 and Md. An1 and An2 each have a three-segmented base (peduncle) and several distal flagellar elements that are not true leg segments because they lack any internal muscles (G. Boxshall & Jaume, 2013; Snodgrass, 1952), meaning that only the peduncle can be compared meaningfully to the other leg types. Additional flagellar elements are added with each molt, and so do not have a set number. An1 is shorter than An2, and An2 has a prominent coxal endite that forms an antennal gland, between which is nestled the labrum.

**Figure 1.**
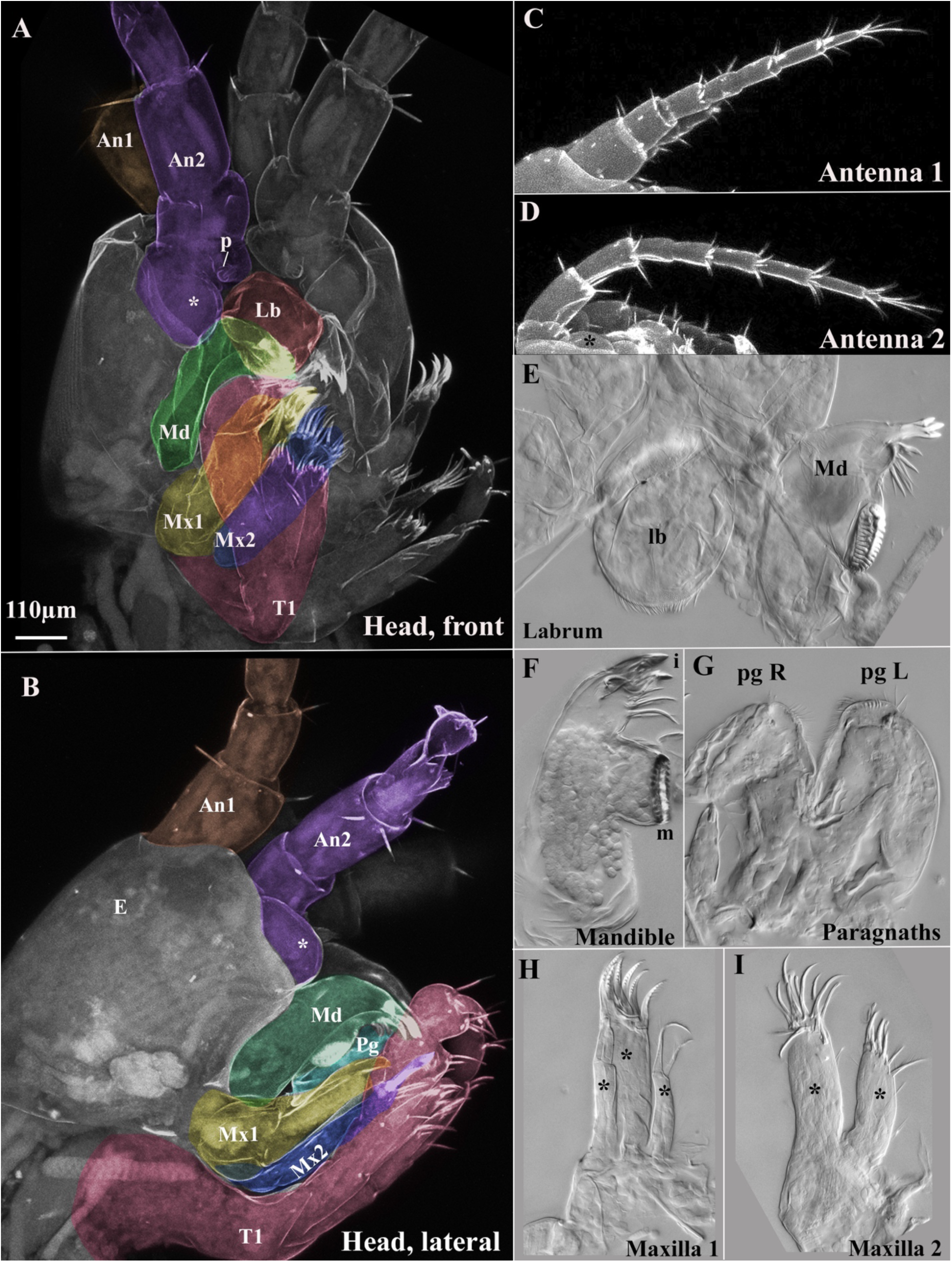
Wild-type morphology of the *Parhyale* head. A. Whole head, frontal view. B. Whole head, lateral view. Antenna 1, brown; antenna 2, purple; labrum, red; mandible, green; maxilla 1, yellow; maxilla 2, cyan; first thoracic leg (T1), magenta. C. Antenna 1 (An1), brown. D. Antenna 2 (An2) with antennal gland coxal endite indicated with asterisk (*), purple. E. Dissected labrum (lb) with mandible (Md). F. Dissected mandible (Md). G. Dissected paragnaths (pg) of the mandible. H. Dissected maxilla 1 (Mx1). I. Dissected maxilla 2 (Mx2). Endites are indicated with an asterisk (*); p, antennal pore.

**Figure 2.**
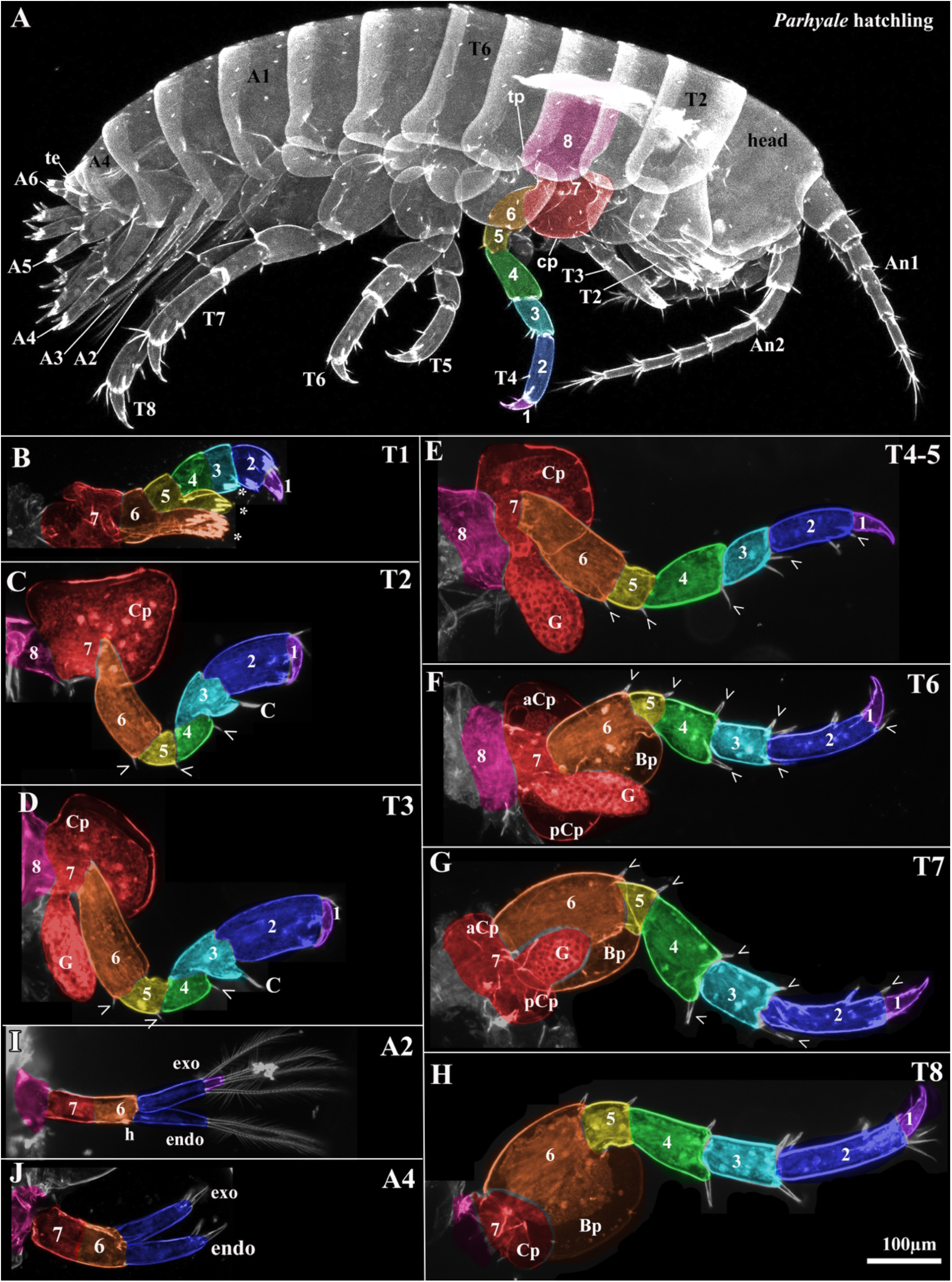
Wild-type morphology of the *Parhyale* thorax and abdomen. T1-8, thoracic body segments 1-8 with corresponding legs. A1-6, abdominal body segments 1-6 with corresponding legs. A. Whole hatchling. B. Dissected T1. C. Dissected T2. D. Dissected T3. E. Dissected T4 or T5. F. Dissected T6. G. Dissected T7. H. Dissected T8. I. Dissected A2. J. Dissected A4. Dorsal boundary of ls8 (precoxa) is approximate. In dissected legs, posterior is down, proximal is left. *Dac*tyl - leg segment 1, purple; propodus - leg segment 2, blue; carpus - leg segment 3, cyan; merus - leg segment 4 green; ischium - leg segment 5 yellow; basis/basipod - leg segment 6 orange; coxa - leg segment 7, red; precoxa - leg segment 8, magenta.

Arthropod mouthparts are generally considered to correspond to the leg base (“gnathobasic”)(G. A. Boxshall, 2004; Popadić et al., 1998; Scholtz et al., 1998), to lack a telopod, which corresponds to leg segments 1 – 5 (ls1 – 5); and are equipped with medial lobes called endites (Fig. 1). The arthropod mandible appears to consist of a single, bilobed endite(Coulcher & Telford, 2013) that forms the incisor and molar with bristles near the incisor. Unlike the mandibles of some other crustaceans and insects, the mandible of *Parhyale* does not have a palp (telopod, or leg segments 1 – 5). The paragnaths are lobes that arise from the medial base of each mandible (Wolff & Scholtz, 2006) and fuse together to form a “lower lip”. Mx1 has three endites: a broad medial lobe with distinct, birefringent combs that is flanked by two slender lobes, each with two strap-like bristles. Mx2 has two nearly-symmetrical endites, one slightly smaller than the other, and each endite topped with a mass of stout bristles on the distal tip.

In the thorax of amphipods (Fig. 2), the first thoracic segment (T1) is incorporated into the head as a maxilliped and functions with the mouthparts. T1 is shaped into a minute claw, but ls5 and ls6 (ischium and basis) each have an endite, the ls5 endite being shorter than the ls6 endite, such that both extend to the middle of ls3 (carpus). The left and right ls7 (coxa) T1 are fused together. T2 and T3 are claws (chelipeds), each with a long, feathery comb on ls3 (carpus).; ls2 (propodus) of T3 is longer than that of T2. T4 and T5 are forward-facing, slender walking legs with no obvious differences between them. T6 – 8 are backwards-facing jumping legs, with T6 being the smallest and stouter, T7 being larger, and T8 being the largest. Each thoracic leg has a bristle on the distal end of each leg segment: in T2 – 5, these distal bristles occur only on the posterior of each leg, while in T6 – 8, the distal bristles occur on both the anterior and posterior leg.

The tergal, coxal, and basal plates on ls8, ls7, and ls6, respectively, cover and protect the animal (Fig. 2). These plates are a type of exite: a lateral lobe on the leg such as a gill or plate that is composed of a bilayer of ectoderm and which may serve a variety of functions (G. A. Boxshall & Jaume, 2009). Each thoracic leg has a tergal plate (tp) emerging from ls8 (the precoxa that is partially incorporated into and now functions as the lateral body wall (Bruce & Patel, 2020)). In T1, where the leg base is fused with the head mass, the tergal plate forms a flange on the head, perhaps together with tergal plates of other head appendages (Bruce & Patel, 2020, 2022; Clark-Hachtel & Tomoyasu, 2020, p. Fig. 1j). The tergal plates project posteriorly and progress from a more rounded shape in anterior legs to more pointed in posterior legs, with T2 having an additional anteriorly-projecting tergal plate that covers the flexible division between the head mass and T2. The coxal plates (cp) project anteriorly and/or posteriorly (Fig. 2). T2-5 have anterior coxal plate lobes, while T6-8 have bilobed coxal plates with both an anterior and a posterior lobe: T6 lobes are equally sized, T7 has a small anterior and large posterior lobe, and T8 has a tiny anterior and large posterior coxal plates. The T2 coxal plate has a distinctive triangular shape, those of T3 – T5 are square shaped, and that of T6 is butterfly-shaped. In T6 – 8, ls6 is expanded into a flat posterior plate (basal plate, bp) that extends over ls5 (Fig. 2). In addition, thoracic legs T3 – 7 each have a gill on ls7, while T2 and T8 do not have a gill. Thus, each thoracic leg is readily identifiable by multiple morphological characters. The only legs that cannot be readily distinguished from each other are T4 and T5.

The abdomen bears biramous appendages where the main axis of the limb is split into a lateral exopod and a medial endopod (Fig. 2). Exopods are different from exites in that exopods have muscle insertions and are often segmented, with muscles that insert on the segmental joint (G. A. Boxshall, 2004; G. A. Boxshall & Jaume, 2009) whereas exites lack muscle insertions and true segmentation. Abdominal legs A1 – 3 are biramous swimmerets, while A4 – 6 are biramous anchoring uropods, and the posterior terminus adjacent to the proctodeum bears tiny, paired appendages of unknown affinity called the telson. The A1 – 3 swimmerets become sequentially slightly smaller, and each exopod is two-segmented with 4 feathery setae (hair-like bristles), and each endopod is one-segmented with 2 setae. There is a patch of tiny Velcro-like hooks on the medial base of each swimmeret that keep the left and right swimmerets together for efficient swimming. The A4 – 6 uropods are smaller and stouter than the swimmerets; they lack the long feathery setae and instead have many stout spines, and become sequentially smaller, such that the A6 uropods are as small as the telson.

## RESULTS

In the early development of *Parhyale*, each cell at the 4-cell stage gives rise to roughly one quadrant of the animal (Browne et al., 2005; Gerberding, 2002). Thus, while CRISPR-Cas9 results in mosaic animals, the affected regions are generally large and contiguous. The following phenotypes are taken from animals where the knockout appears to have occurred early in development and affects all or large quadrants of the animal.

## FUNCTION OF DISTALLESS IN *PARHYALE*

*Dll* CRISPR-Cas9 knockout truncates all legs, leaving only ls6 – 8 (Fig. 3). An1 and An2 are truncated. In An1, three peduncle segments remain, along with one or two distal elements that may be flagellar in nature. In An2, two peduncle segments remain: the peduncle segment with the coxal endite, another peduncle segment, and one distal element that may be flagellar in nature. Notably, the shortened antennae do not terminate in an “unfinished” stump like the other legs, but rather the distal (flagellar?) tip is always pointed and tipped with bristles. Although *Dll* is expressed in the mandible, mx1, and mx2 (Fig. 4), these limbs do not appear to be affected by loss of *Dll* (Fig. 3), even when adjacent structures, like the labrum, antennae, and T1, are severely affected, suggesting that *Dll* was indeed knocked out in these tissues, and that the lack of phenotype in the mouthparts is not because they are wild-type. *Dll* expression in the mouthparts may be patterning sensory structures like bristles. Notably, *Dll* is expressed in the tip of the mandible, even though no palp is present in amphipods; this *Dll* domain likely represents the location where the palp develops in other mandibulates. In the A1 – 3 swimmerets and A4 – 6 uropods, the exopod and endopod are deleted, but usually a long, feathery seta or two are still present. Interestingly, in both swimmerets and uropods, the endopod is more reduced than the exopod. This could mean that the endopod is more sensitive to the loss of *Dll*, or that the exopod requires less *Dll*. The telson is not affected. The *Dll* expressing bristle on posterior distal ls6 is usually deleted, but not the anterior distal bristle. Despite the expression of *Dll* in the tergal plate, coxal plate and gill (Bruce & Patel, 2020) (Fig. S1), these structures appear wild-type in *Dll* knockouts.

**Figure 3.**
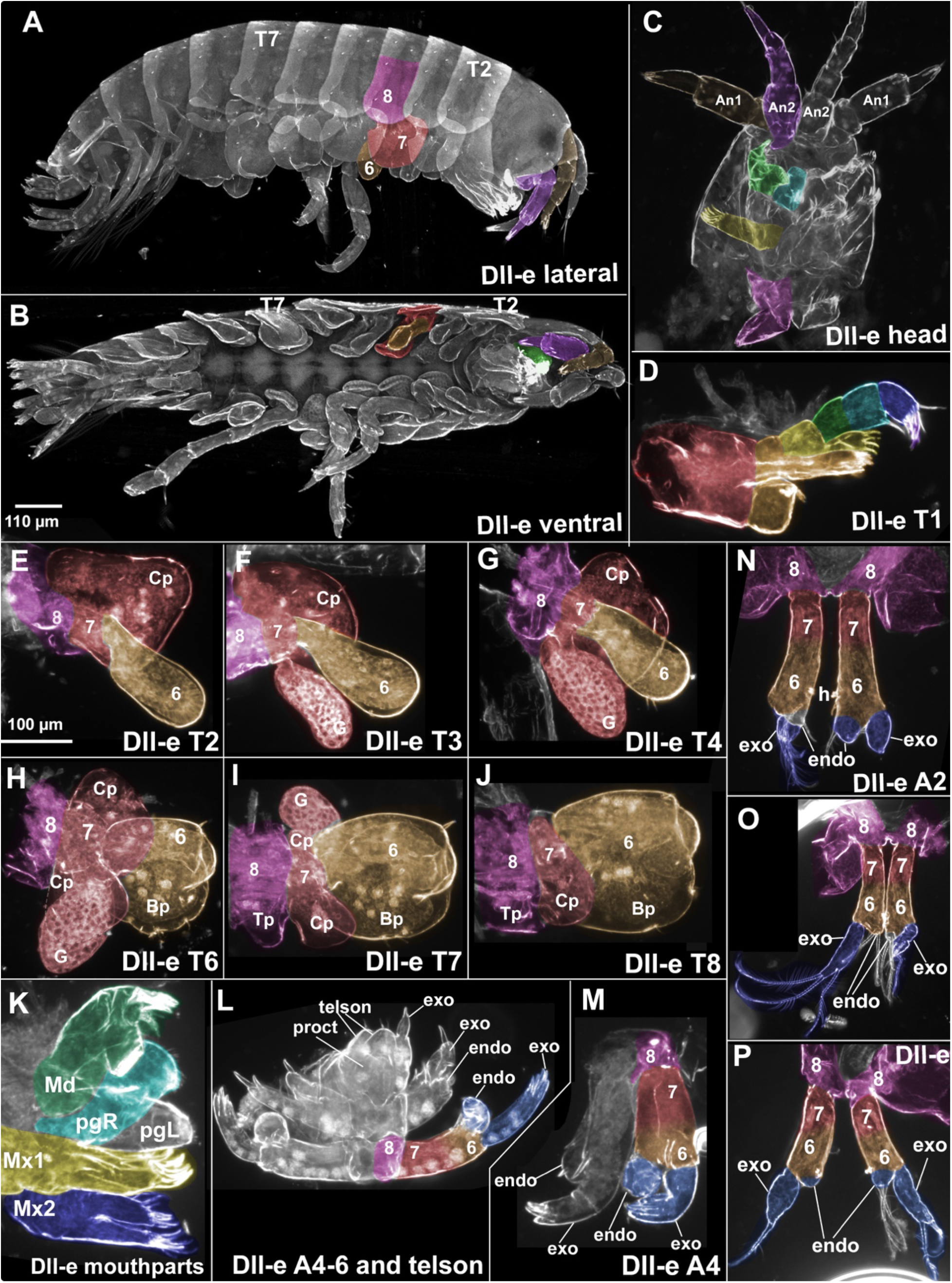
Distal-less phenotypes in *Parhyale* appendages. Leg segments 1-5 deleted. Colors and numbers in head as in Figure 1 legend, colors and numbers in thoracic and abdominal leg segments as in Fig. 2 legend. A, whole hatchling, lateral view. B, whole hatchling ventral view. Antenna 1 (An1) in brown, antenna 2 (An2) in purple, remaining leg segment of T2 leg colored as in Figure 2 legend. C, dissected head. Labrum is deleted, and An1, An2, and T1 severely affected, but mouthparts are unaffected. D, Dissected T1. E, Dissected T2. F, Dissected T3. G, Dissected T4 or T5. H, Dissected T6. I, Dissected T7. J, Dissected T8. Note that in T6-8, the small stump left on ls6 is the distal end of ls6, rather than a portion of ls5. K, dissected mouthparts, mandible (Md), paragnath (pgL, pgR), maxilla 1 (Mx1), maxilla 2 (Mx2). L-P, in all biramous appendages, the endopod is more affected than the exopod, and in some cases, a nearly completely deleted endopod is next to a nearly normal-looking exopod. L, biramous abdominal uropods A4-6 and telson. M, fourth biramous abdominal uropod, A4. N, second biramous abdominal swimmeret, A2. O,P, biramous abdominal swimmerets. Dorsal boundary of ls8 (precoxa) is approximate. In dissected thoracic legs, posterior is down, proximal is left..

## FUNCTION OF SP6-9 IN PARHYALE

*Sp6-9* CRISPR-Cas9 knockout affects ls1 – 6 (Fig. 3). An1 and An2 are truncated and shorter than in *Dll* KOs, leaving the segment that carries the coxal endite, along with one or two distal elements that may be flagellar in nature. The labrum is smaller but not entirely deleted as it is in *Dll* knockouts. The tergal plate, coxal plate, and gill are unaffected and appear wild-type.

Neither the mandible nor its paragnath are affected, even when adjacent structures, like antennae and T1, are severely affected. In Mx1, one of the small endites is deleted. In Mx2, the smaller endite is withered or leaves only a swelling.

The phenotype in T1 is difficult to interpret: it deletes ls1 – 6 as expected, but in addition, it leaves what looks like a normal endite emerging from ls7, even though no endite exists on ls7 of T1 in WT animals. In T2 – 5, the small nub emerging from the coxal plate is the ls7 joint that normally connects to ls6. In T6 – 8, a smaller, misshapen ls6 is still present. This may be related to the weaker *Sp6-9* expression in proximal ls6 of these legs (Fig. 6). The swimmerets and uropods are truncated and no remnants of the exopods or endopods remain, but part of ls6 appears to remain. The telson is not affected.

Another interesting *Sp6-9* phenotype in T6 – 8 is that the basal plate on ls6 often narrows where it joins the leg segment. This is similar to the shape of the coxal plate on ls7, which is also narrow where it joins the leg segment. This phenotype suggests that loss of *Sp6-9* weakly transforms ls6 of T6 – 8 towards a coxal identity. This is further supported by the phenotype in T6, the only leg where the WT coxal plate is bilobed with an anterior and posterior lobe. In T6 *Sp6-9* KO hatchlings, the basal plate is also becoming bilobed with an anterior and posterior lobe. Intriguingly, loss of *Sp6-9* in *Drosophila* also occasionally led to transformations of distal leg regions into more proximal ones (Estella & Mann, 2010): the distal leg was sometimes transformed into notum (ls8, precoxa) with wing tissue (exite). Another interesting phenotype in T6 – 8 is that the anterior distal edge of ls6 (“basipod”) often forms a nub with clusters of bristles and stout spines, which could possibly represent a weak transformation towards a biramous limb like the uropods (Figs. 4, S1).

**Figure 4.**
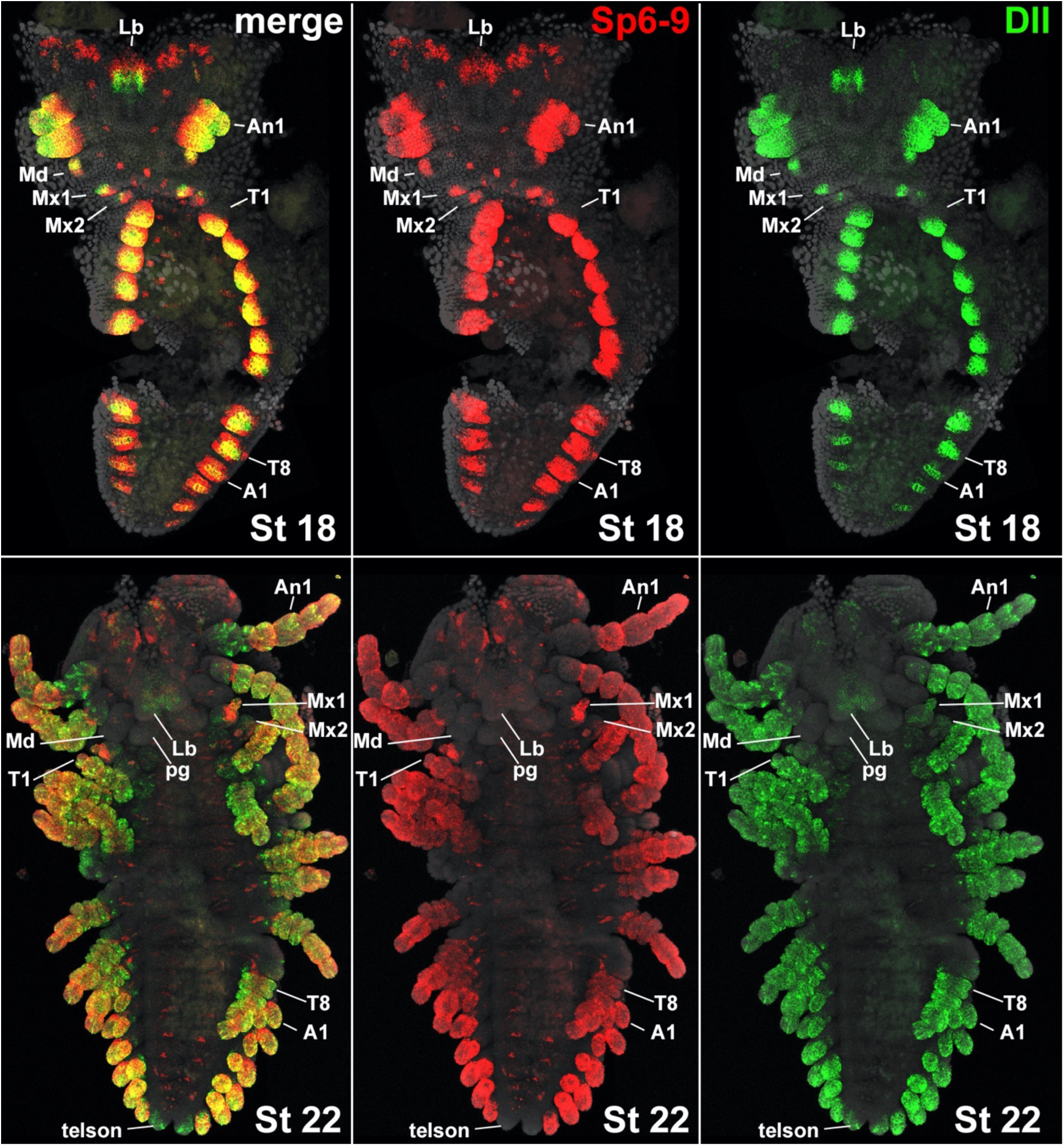
Expression of *Dll* and *Sp6-9* in Stage 18 and 22 whole *Parhyale* embryos. At St18, *Sp6-9* expression surrounds that of *Dll*, consistent with a more proximal patterning function. Both are expressed in the tip of the mandible as well as Mx1 and Mx2. In the labrum, *Sp6-9* is expressed in a more dorsal position. *Sp6-9* is expressed in a few cells medial to each thoracic leg, and in a smattering of cells around the head lobes. At St22, *Dll* is expressed in streaks in An1, which may represent a sensory function, and *Dll* is expressed in the telson. At St22, the *Sp6-9* expression medial to each leg appears to now form muscles, while some of the *Sp6-9*-expressing head lobe cells now reside in a pair of kidney-shaped thickenings (ocelli?) dorsal to the labrum, and others are arrayed around the ocular segment (somewhat folded behind the rest of the head).

Notably, the distal end of the truncated limb is often ballooned out (Fig. 7 and Fig. S1). This occurs most frequently in the antennae, where the distal tip can balloon to half the size of the head. In both mx1 and mx2, the distal ends are frequently ballooned, or end in numerous small blebs (Fig. S1). The distal swimmerets are occasionally ballooned as well.

## FUNCTION OF *DAC*HSHUND IN *PARHYALE*

*Parhyale* has two copies of *dac*, *dac1* and *dac2*. Only *dac*2 gives a phenotype, but *dac1* and *dac2* have similar but not identical expression patterns (Fig. S2). *Dac2* CRISPR-Cas9 knockout deletes ls3 – 5 (Fig. 5). Only the thoracic legs appear to be affected, as the appendages of the head (An1, An2, Lb, Md, pg, Mx1, Mx2), and abdomen (swimmerets A1 – 3 and uropods A4 – 6) have the WT number and size of elements. The telson is not affected. In the thoracic legs of the most severely affected hatchlings, ls3 – 5 are completely deleted such that ls2 and ls6 connect directly to each other and share a joint, while in less severely affected hatchlings, ls2 and ls6 are separated by a short ring of tissue that represents the fused ls3 – 5 (Fig4.11). ls5 was only affected in the most severely affected animals, and ls5 also has a low expression level of *dac*2.

**Figure 5.**
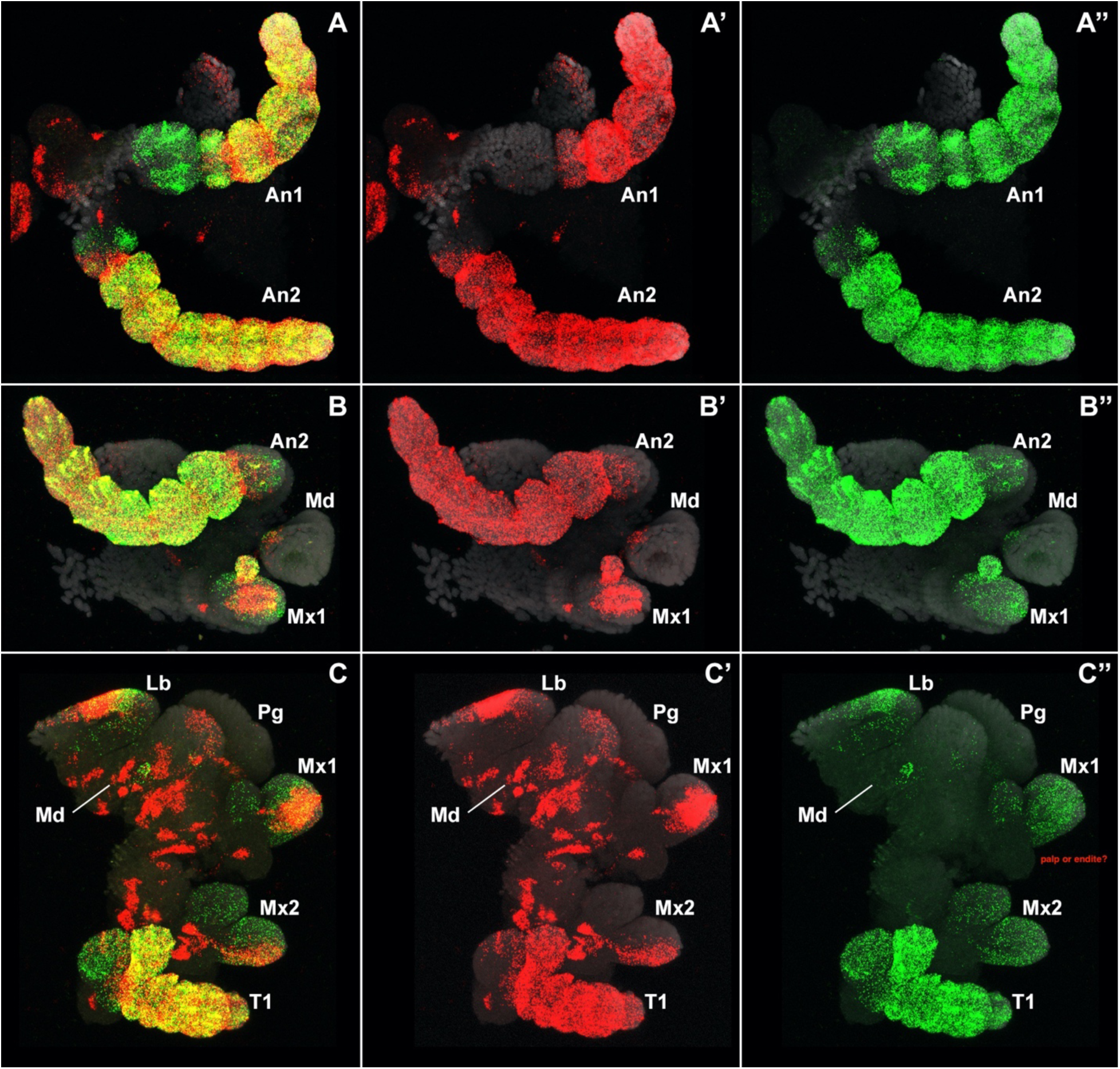
Expression of *Dll* and *Sp6-9* in *Parhyale* dissected head and mouthparts. In the labrum, *Sp6-9* is expressed in a more proximal domain than *Dll*, and both are expressed in speckles throughout the labrum. In An1 and An2, *Dll* is expressed in streaks and extends more proximally than *Sp6-9*, both of which perhaps reflect a sensory function for *Dll* here. *Dll* and *Sp6-9* are expressed in the tip of the mandible as well as Mx1 and Mx2. *Dll* is expressed in a more proximal domain than *Sp6-9* in Mx1, Mx2, and T1, even though this region is not deleted when *Dll* is knocked out, which suggests a sensory function for *Dll* here. The streaks of *Sp6-9* expression in the mouthparts are likely serially homologous to streaks of *Sp6-9* medial to each thoracic leg, and likely mark muscles.

Conversely, *dac* expression is highest in ls4, and the mildest *dac* phenotype causes ls4 to be truncated and fused onto ls3. Interestingly, in several severely affected hatchlings, there was a dorsal nob on ls2 of several thoracic legs. In T1, ls3 – 5 are reduced, as they are in the other thoracic legs, and the endite on ls5 is absent. This becomes apparent when comparing the longer endite of the unaffected ls6 to the main axis of the leg: normally the ls6 endite ends in the middle of ls3, but when *dac*2 is knocked out, the ls6 endite extends a full segment further at ls2. Perhaps the partial maxillary identity of T1 that is conferred by Scr (Abzhanov & Kaufman, 1999; Liubicich et al., 2009; Martin et al., 2016; Pavlopoulos et al., 2009; Serano et al., 2015) buffers the medial leg segments of T1 against complete deletion.

## FUNCTION OF EXTRADENTICLE AND HOMOTHORAX IN *PARHYALE*

*Exd* and *Hth* in *Drosophila* have been shown to act as cofactors that require co-expression to be transported into the nucleus and to prevent their degradation, and as such, their knockout phenotypes are nearly identical (Mito et al., 2008; Pai et al., 1998; Ronco et al., 2008). Knockout phenotypes in *Parhyale* also give apparently identical phenotypes (Figs. 6, 7).

**Figure 6.**
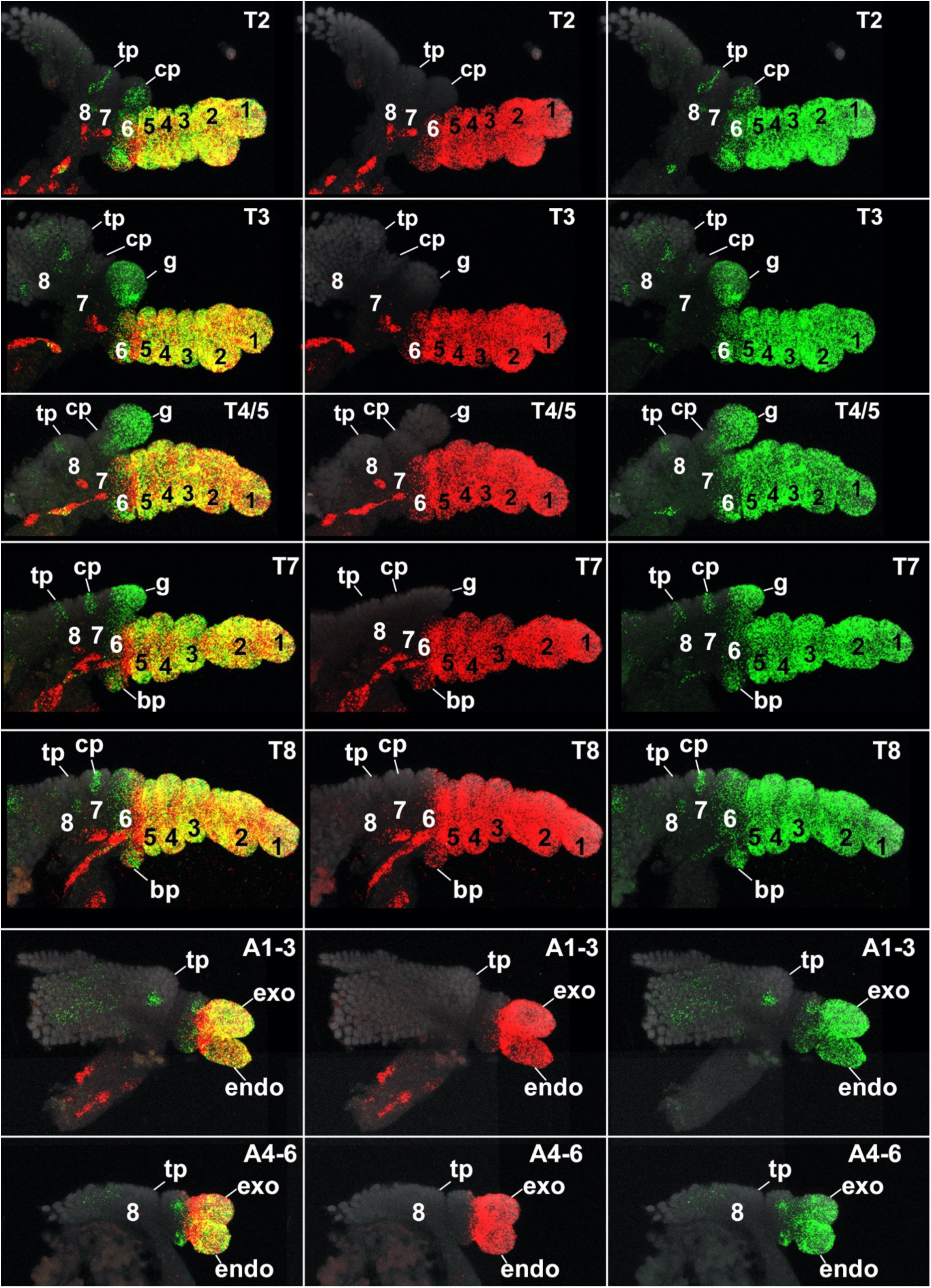
Expression of *Dll* and *Sp6-9* in *Parhyale* dissected thoracic and abdominal limbs. *Dll* is expressed in all exites: in a small streak in the tergal, coxal, and basal plates, and throughout the distal tip of each gill (not just on the superficial surface of the gill). This difference in expression may represent a means to distinguish exites of unknown function in other arthropods. *Sp6-9* is expressed in ventral streaks that appear to be muscles. In the biramous abdominal swimmerets and uropods, *Dll* extends more proximally than *Sp6-9*.

**Figure 7.**
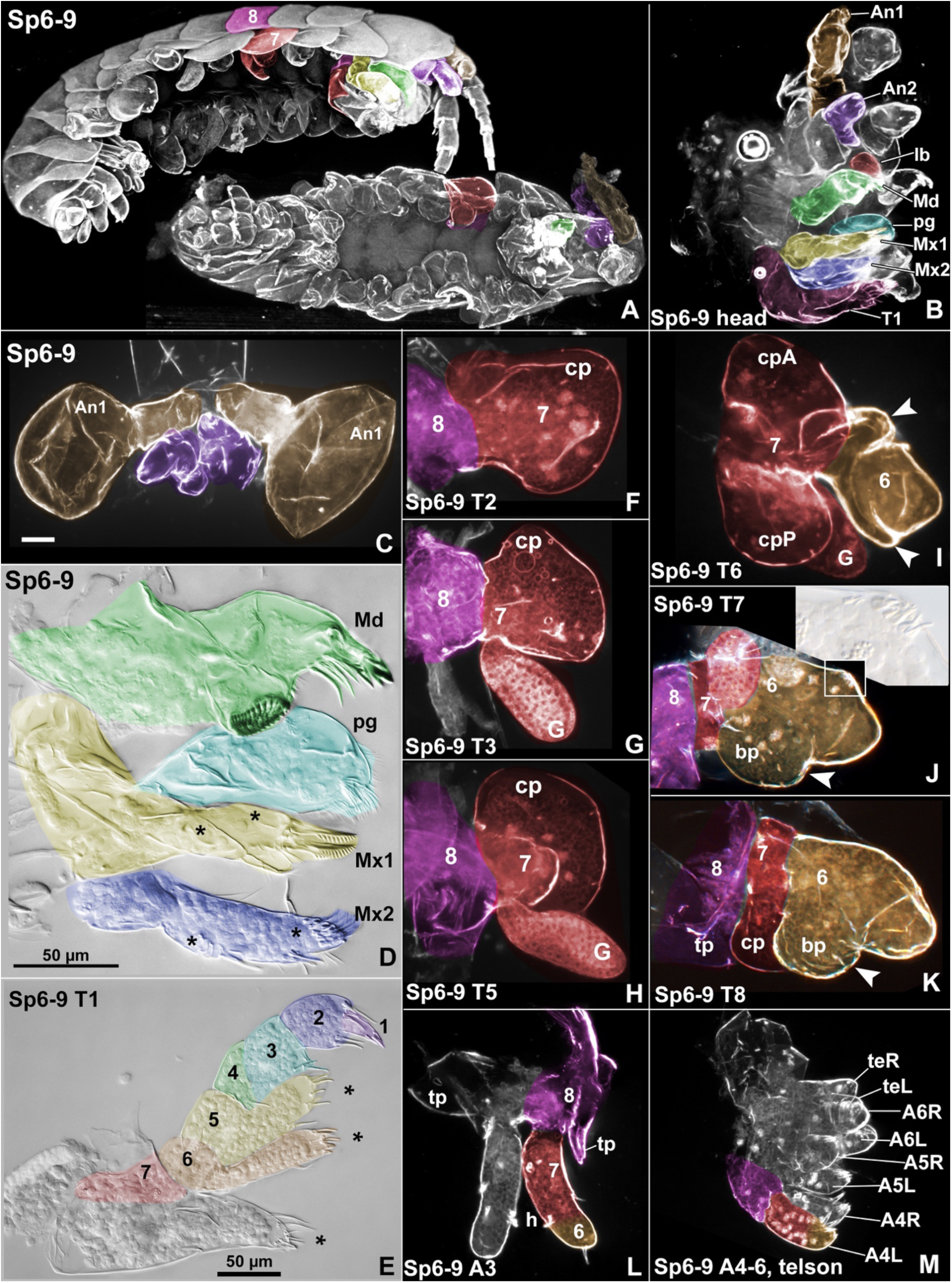
*Sp6-9* phenotypes in *Parhyale*. Leg segments 1-6 deleted. Colors and numbers in head as in Figure 1 legend, colors and numbers in thoracic and abdominal leg segments as in Fig. 2 legend. Dorsal boundary of ls8 (precoxa) is approximate. Endites marked with asterisk (*). A, Oblique and ventral views of whole *Parhyale* hatchlings. B, Dissected head. C, Dissected An1 with ballooned distal tissue, and An2. D, Dissected mandible (Md), paragnath (pg), maxilla 1 (Mx1), and maxilla 2 (Mx2). Md is unaffected. E, thoracic limb 1 (T1). F-K, thoracic legs T2-8. Closed arrowhead marks the neck of the basal plate (bp) in J and K, and bp transformed towards coxal plate identity in I. Inset in J shows ectopic lobe with stout spines. L, third biramous abdominal swimmeret, A3. M, biramous abdominal uropods A4-6 and telson, teR and teL. Tergal plate, tp; coxal plate, cp; gill, g.

The strongest phenotypes when either *Exd* or *Hth* is knocked out results in fusion and deletion of proximal leg segments leaving distal leg segments 1, 2, and distal part of 3 intact (Figs. 6, 7). Deletion of proximal leg segments includes the lateral body wall with its tergal plate, which together are derived from the embryonic precoxa or ls8 (Bruce, 2022; Bruce & Patel, 2020). In the most severely affected hatchlings, multiple normal, distal legs emerge from a single smooth region of body wall. The labrum and surrounding tissue are affected: the peripheral edge of the labrum is reduced such that the labrum is pointed rather than mound-shaped. The telson is not affected.

Notably, the mandibles are never affected despite being a proximal leg segment. This is likely due to the lack of *Hth* expression in this appendage (Fig. S3). Similarly, the A1 – 3 swimmerets and A4 – 6 uropods are also normal (Figs. 6, 7), except for the lateral body wall (precoxa or ls8) from which they emerge, which is often fused and distorted. The lack of a phenotype in these biramous appendages is not likely a result of mosaicism with WT tissue because the adjacent tissue is strongly affected - normal-looking swimmerets and uropods emerge from fused lateral body, and the proximal leg segments of the adjacent T8 leg are fused and deleted. Moreover, *exd* appears to be weakly expressed throughout the abdomen (Fig. S3).

*Exd* or *Hth* CRISPR-Cas9 knockout animals often develop a contiguous, fused head and thorax (Fig. 6, 7). These fusions/deletions are more frequent and severe in the anterior body region. Unilateral fusions that are restricted to the proximal leg (i.e. tergum or lateral body wall) of what appears to be T2-4 occur frequently. Hatchlings with such unilateral fusions and deletions generally develop to hatching. The foreshortening of only the left or right half of the body twists the hatchling into a nearly spiral shape. Hatchlings with bilateral fusions/deletions can’t hatch out by themselves and must be hand dissected, and the cuticles do not have time to harden, resulting in wrinkly cuticles.

More moderate phenotypes result in transformations of leg types. Mouthparts on the head lose their endites and develop into long but generic uniramous appendages. Thoracic legs develop both mouth-like endites and abdomen-like exopods. T2 and T3 are transformed towards a generic thoracic leg identity and the thick claw of ls2 becomes slender. In milder *exd* or *hth* phenotypes, discrete regions of red pigment form on the antennae, T1, and pharynx (Fig. S1), which appear to be ectopic eye tissue.

## DISCUSSION

*Dll* knockout in *Parhyale* deletes leg segments 1 – 5 (*dac*tyl – ischium). Notably, *Dll* KO does not alter the morphology of the tergal plate, coxal plate, basal plate, or gill, even though *Dll* is expressed in these exites. This mirrors the situation in the insect wing, which has been shown to be a type of leg exite that is positionally homologous to the crustacean tergal plate (Bruce & Patel, 2020). In *Drosophila*, *Dll* is strongly expressed in the wing margin, but loss of *Dll* does not affect the overall morphology of the wing blade (Gorfinkiel et al., 1997). Instead, *Dll* expression in the wing margin appears to have a sensory function, as loss of *Dll* deletes only the sensory bristles here (Panganiban, 2000). In *Parhyale*, *Dll* also appears to pattern bristles on exites. Given that *Dll* regulates the proneural gene *achaete-scute* in the *Drosophila* wing (Campbell & Tomlinson, 1998; Panganiban, 2000), a sensory function for *Dll* in the *Parhyale* gill and coxal plate could be assayed by examining *achaete-scute* expression in *Parhyale* exite bristles.

*Sp6-9* knockout in *Parhyale* deletes ls1 - 6 (*dac*tyl – basis). The abrupt difference in the degree of deletion of ls6 between T5, where ls6 is completely deleted, and T6, where a large portion of ls6 is intact, suggests the influence of a Hox gene. AbdA and abd-B are likely candidates, as these are expressed in the posterior of the animal starting with T6. If insects and spiders had full length legs on their posterior trunk like crustaceans do, then *Sp6-9* function might similarly be reduced in ls6 of these posterior trunk legs. This theory could be tested in a myriapod (millipedes, centipedes, etc.) However, the anterior trunk legs of all arthropods follows the same pattern for *Sp6-9*, wherein ls6 is completely deleted.

Notably, exites (gills, plates, etc) express *Dll* but not *Sp6-9*. This difference can be exploited to determine the identity of ambiguous lateral outgrowths in arthropods. If the outgrowth is an exite, it will express *Dll* but not *Sp6-9*. If it is an exopod, it will express both *Dll* and *Sp6-9*, with *Sp6-9* expressed more proximally than *Dll*.

*Exd* or *Hth* knockout phenotypes were qualitatively indistinguishable from each other.

Transformations of one leg type into another frequently occur in animals moderately affected by *exd* or *hth* knockout. The Hox genes confer different leg identities, such as claw, walking leg, or swimmeret, and *Exd* and *Hth* are well-known cofactors of Hox genes (Martin et al., 2016; Slattery et al., 2011). Thus, when *Exd* or *Hth* levels are moderately reduced – enough to affect interactions with the Hox proteins, but not so low as to delete the proximal leg segments – this results in transformations in identity from one leg type toward another. Previously, these transformations have been interpreted in insects and spiders as transformations of head appendages into thoracic appendages(Casares & Mann, 2001; Inbal et al., 2001; Pai et al., 1998; Pichaud & Casares, 2000; Rieckhof et al., 1997; Sharma et al., 2015). However, these interpretations of *exd* and *hth* behavior may be constrained by the limited diversity of appendages in insects and spiders. In contrast, in a crustacean like *Parhyale* – with twelve or more distinguishable types of legs on the head, thorax, and abdomen – a more fine-grained assessment can be made between various leg identities. In *Parhyale*, loss of *exd* or *hth* in the head appendages causes transformation towards a thoracic (uniramous) identity (Figs. 6, 7), but in the *Parhyale* thorax, loss of *exd* or *hth* seems to lead, not to a unidirectional transformation of head legs into thoracic legs, but to a general permissiveness to form various leg components (Fig. 6). For example, following the loss of *exd* in *Parhyale*, endites - normally found only on the head - and biramous appendages - normally found only on the abdomen - can be found together in each leg of the thorax. This means that a single thoracic leg displays a mosaic of characters from head, thorax, and abdomen. A similarly permissive effect may also be happening in insects and spiders, but may be less obvious simply due to the paucity of leg components in these arthropods. Thus, many structures may be competent to form in most body regions, but the Hox genes appear to be suppressing specific structures to only certain body regions. Presumably, these suppressed structures could be allowed to form under the right circumstances, and may be a source of new structures and new functions in evolution.

*Exd* and *hth* are generally thought to pattern the body wall in addition to the proximal leg.

However, given that the lateral body wall of many arthropods is derived from the incorporated leg base (Bruce, 2022; Bruce & Patel, 2020; Kobayashi et al., 2022), the lateral “body wall” patterned by *exd* and *hth* likely corresponds to this incorporated leg base, meaning that *exd* and *hth* probably do not pattern the dorsal true body wall. This is also supported by the expression of *exd* in onychophorans, a lineage which branched prior to the evolution of arthropod leg segmentation (“arthropodization”) and thus prior to the incorporation of proximal leg into the body wall in various arthropods. In onychophorans, *exd* expression forms a tight ring around the base of each leg and does not extend into the trunk body wall (Janssen et al., 2010). Rather than *exd* or *hth*, *pannier* (*pnr*) seems to pattern the true body wall that is not derived from the ancestral/embryonic leg.

Based on the phenotypes presented here, the mouthparts - Md, Mx1, and Mx2 - appear to correspond to proximal leg segments, which confirms previous interpretations of mandibulate arthropod mouthparts as being gnathobasic. The expression of *araucan* (*ara*, Fig. S3), has been shown to bracket the ls8 (precoxa) in all arthropods examined (Bruce, 2022; Bruce & Patel, 2020, 2022, p. 202; Diez del Corral et al., 1999). Based on *ara* expression in the head, the region of the lateral head mass from which the mouthparts emerge is bracketed dorsally and ventrally by *ara*, and thus corresponds to ls8 (precoxal) tissue (compare similar expression of *ara* in the head to that in the thorax and abdomen). This smooth, contiguous head cuticle thus appears to consist of a conglomeration of multiple ls8’s (precoxae), one from each of the head appendages (An1, An2, Md, Mx1, Mx2, and T1). The lack of *Dll* phenotype in the mouthparts suggest that ls1 - 5 (*dac*tyl - ischium) are not present in these legs. Since ls8 (precoxae) forms the lateral head mass (based on *ara* expression) and ls1 - 5 are not present (lack of *Dll* phenotype), this molecular triangulation method means that the free and mobile parts of the mouthparts must consist of only two leg segments: ls6 and ls7 (basis and coxa). The regions of the mouthparts affected by *Sp6-9* knockout should correspond to ls6 (basis), while the remaining unaffected regions should correspond to ls7 (coxa). Furthermore, the ls8 (precoxa) of each mouthpart that forms the lateral head mass presumably continues medially. Given that the paragnaths resemble and function similar to endites, and emerge from the mandibular “sternum”(Wolff & Scholtz, 2006) – here proposed to be the precoxa (ls8) – the paragnaths may represent the endites of the Md precoxa.

In An1 and An2, *Dll* and *Sp6-9* appear to be involved in patterning both the peduncle segments that correspond to true leg segments, and the flagellar subdivisions that do not. *exd* and *hth* appear to affect the peduncle but not the flagellum, while *Dac* does not seem to affect the antennae. The stripe of *ara* at the base of ls7 (coxa) of An1 and An2 means that ls8 (precoxa) of each antennae forms part of the lateral head capsule. The streak of *ara* in the mobile region of An2 likely corresponds to ls6 (basis), as it does in the other legs, so the interpretation of the antennal gland as a coxal endite is consistent with the molecular data presented here. The peduncle seems to correspond to the base of the leg of the post-antennal legs, while the flagellum - with its lack of intrinsic muscles and absence of distinctive, individuated elements - seems to be an alternative distal program that is unrelated to the distal region of the post-antennal legs.

The identity and affinities of the arthropod labrum - and whether it is in some way related to the canonical legs - has been a vexing problem for generations of arthropod researchers (see “arthropod head problem” and (Budd, 2002; Ortega-Hern ndez et al., 2017; Panganiban et al., 1994; Scholtz & Edgecombe, 2006). Like canonical legs, the labrum develops as paired nubs from a pair of coelomic sacs and is patterned by essentially the same gene network, including the leg genes examined here (Abzhanov & Kaufman, 2000; Bruce & Patel, 2020; Panganiban et al., 1994; Pechmann & Prpic, 2009; Prpic et al., 2001; Prpic & Tautz, 2003; Scholtz & Edgecombe, 2006) and segmentation genes like wingless (Janssen et al., 2008; Murat et al., 2010; Prpic & Tautz, 2003). However, no convincing transformation of the labrum into a canonical leg has been observed, as would be expected for a serial homolog (Haas et al., 2001). The results reported here for *Parhyale* are similarly ambiguous: the *Parhyale* labrum is affected by *Dll*, *Sp6-9*, *exd*, and *hth*, but not *dac*. While this is consistent with some kind of leg affinity, the phenotypes of these leg patterning genes in the labrum are not consistent with their phenotypes in canonical legs. For example, *Dll* completely deletes the labrum, but *Sp6-9* only reduces it, whereas in no canonical leg was the *Dll* phenotype more severe or more proximal than the *Sp6-9* phenotype. Furthermore, while loss of *exd* or *hth* deleted what could be interpreted as the “proximal” labrum (peripheral edge of the labrum), there were no transformations of the labrum into a canonical leg type. One possible explanation for these conflicting data is that the labrum may be derived from an ancient proto-limb, from which both the labrum and trunk limbs later evolved (Fig. 8). The majority of the bilaterian body today is patterned by the Hox genes, but there is a small anterior region of the head where Hox genes are not expressed, and this anterior Hox-free region is instead patterned by other homeobox-family genes like orthodenticle (ocelli-less) and optix (Posnien 2023; Steinmetz 2010; Marlow 2014). The presence of the An1 limb (chelicera in chelicerates) in this Hox-free anterior region means that this region is capable of forming limbs (and perhaps also that limbs evolved prior to the division of the bilaterian body into a Hox-free anterior and Hox-patterned posterior). Notably, An1 in the Hox-free anterior is indeed serially homologous to the Hox-patterned trunk limbs because An1 can be transformed into a trunk limb with the Antp mutation (Struhl, 1981). It is well-established that body segmentation genes like *wingless* are expressed somewhat differently in the anterior Hox-free region than in the Hox-patterned trunk. This is to be expected given the very ancient divergence of these Hox-free segments from the Hox-patterned segments, which – given that this Hox-free anterior region is patterned by the same genes in the same configuration in both cnidarians and bilaterians (Marlow et al., 2014; Steinmetz, 2010) – likely occurred prior to the divergence of cnidarians and metazoans. The same may be true of the leg patterning genes in the Hox-free anterior region: if an ancient proto-limb gave rise to both the Hox-free labrum and An1 regions, as well as the Hox-patterned canonical legs, and subsequently the Hox-patterned proto-limbs evolved into canonical limbs, then this might explain why so much of the leg-patterning gene network is expressed in the labrum, but in slightly different ways than in canonical legs, and why no convincing transformation of labrum into canonical leg has been achieved. Perhaps mis-expressing the Hox gene Labial or the leg gene Sp6-9 in the labrum could transform the labrum into a canonical leg, thereby demonstrating convincingly that the labrum is a leg serial homolog.

**Figure 8.**
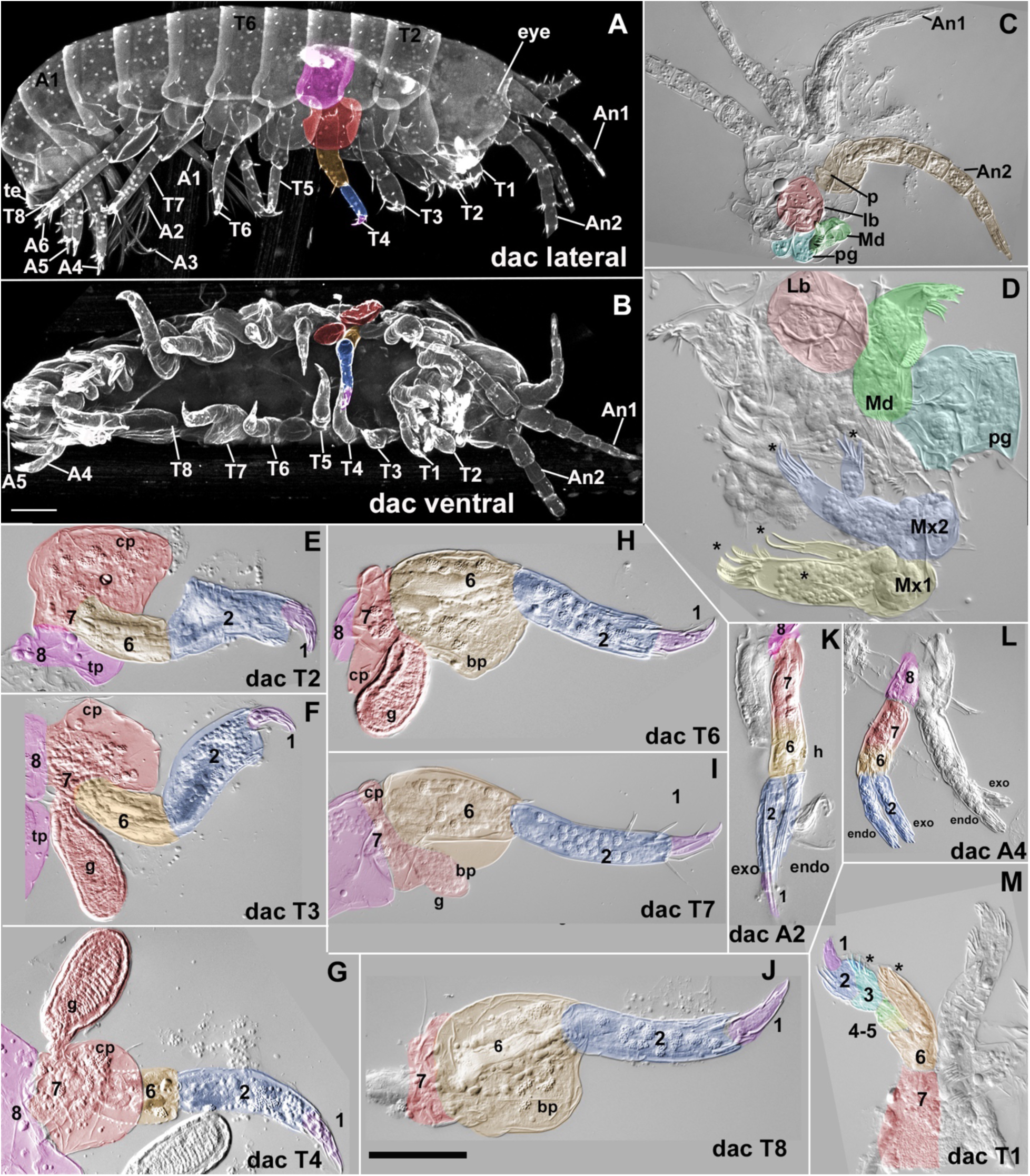
*dac*hshund phenotypes. Leg segments 3-5 deleted. Colors and numbers in head as in Figure 1 legend, colors and numbers in thoracic and abdominal leg segments as in Fig. 2 legend. Dorsal boundary of ls8 (precoxa) is approximate. A, whole hatchling, lateral view. Legs on right side of animal are labeled. Hatchlings could not hatch on their own and had to be dissected from embryo membranes, so the cuticle did not have time to harden before fixation, resulting in delicate, wrinkly cuticle. B, whole hatchling, ventral view. Legs on left side of animal are labeled. Note that the whole mount embryo in ventral view is pressed up against the slide, distorting the swimmerets and setae, but the swimmerets and uropods are WT (compare swimmerets and uropods in ventral and lateral view of whole embryo). C, dissected head. D, dissected mouthparts, labrum (Lb), mandible (Md), maxilla 1 (Mx1), maxilla 2 (Mx2). E-I, dissected thoracic legs T2-8. K, dissected second biramous abdominal swimmeret, A2, size and shape of leg segments is normal. Left swimmeret mostly out of focus behind right swimmeret, except for its distal setae that wrap around to the right swimmeret. L, dissected fourth biramous abdominal uropod, A4, size and shape of leg segments is normal. M, dissected T1, ls3-5 are shortened and endite on ls5 is absent, but other endites (*) are present. ls8 with tergal plate and ls7 with coxal plate accidentally dissected away in some legs.

**Figure 9.**
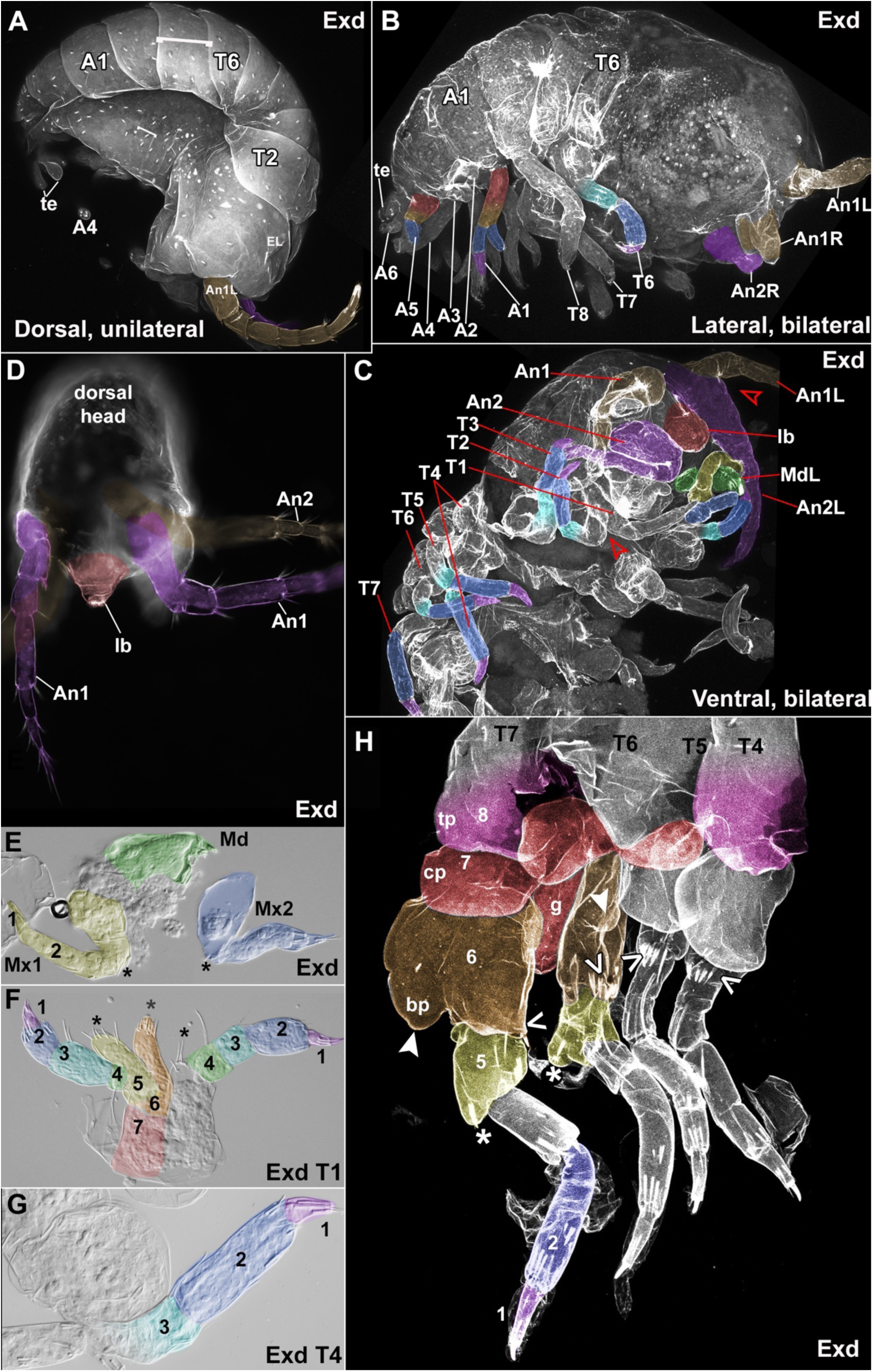
Extradenticle phenotypes. Proximal leg segments deleted, leaving ls1, 2, and the distal part of ls3 normal. Colors and numbers in head as in Figure 1 legend, colors and numbers in thoracic and abdominal leg segments as in Fig. 2 legend. A, whole hatchling, dorsal view, unilateral knockout. A portion of each body segment is deleted, and the remaining body segments are fused together, resulting in a hatchling with a dramatically twisted body. B, whole hatchling lateral. Anterior body is more affected than the posterior. Legs labeled where identifiable. Mobile part of abdominal swimmerets and uropods appear normal, despite emerging from a lateral body wall (=ls8 = precoxa) that is fused and distorted. Note that A2 and A3 appear WT when panning through the z stack, but were accidentally crushed during mounting. Leg segments are fused from proximal ls3, indicated by fading of cyan color. C, whole hatchling, ventral view. Legs labeled where identifiable. Mx1 (yellow), Mx2 (blue), and T1 transformed towards generic uniramous identity. Mandible (green) unaffected. Open arrowheads indicate fusions of adjacent legs at their bases. D, dissected head, doral view. Periphery of labrum is deleted. E, dissected mouthparts, mandible (Md, green), maxilla 1 (Mx1, yellow), maxilla 2 (Mx2, blue). Mx1 and Mx2 transformed towards generic uniramous identity with loss of endites (*) and gain of joints and distal leg segments (ls1 and ls2 discernible in Mx1). Md unaffected. F, T1 left side WT, right side transformed towards generic uniramous identity with loss of endites (*) and gain of larger ls1 and ls2. G, dissected thoracic leg T4. H, dissected thoracic leg T4-7 with transformations towards a mosaic of identities from head, thorax, and abdomen. These thoracic legs bear tergal plates (tp), coxal plates(cp), and basal plates (bp, closed arrowheads), gills (g), and a mostly uniramous identity, but also have ectopic endites (*) on ls5 like T1 (compare to ls5 endite on T1 in G), and exopods from the abdomen (indicated with <, forming lobes with stout spines emerging from the lateral side of ls8, the basis, the location where exopods normally form in the abdominal swimmerets and uropods).

**Figure 10.**
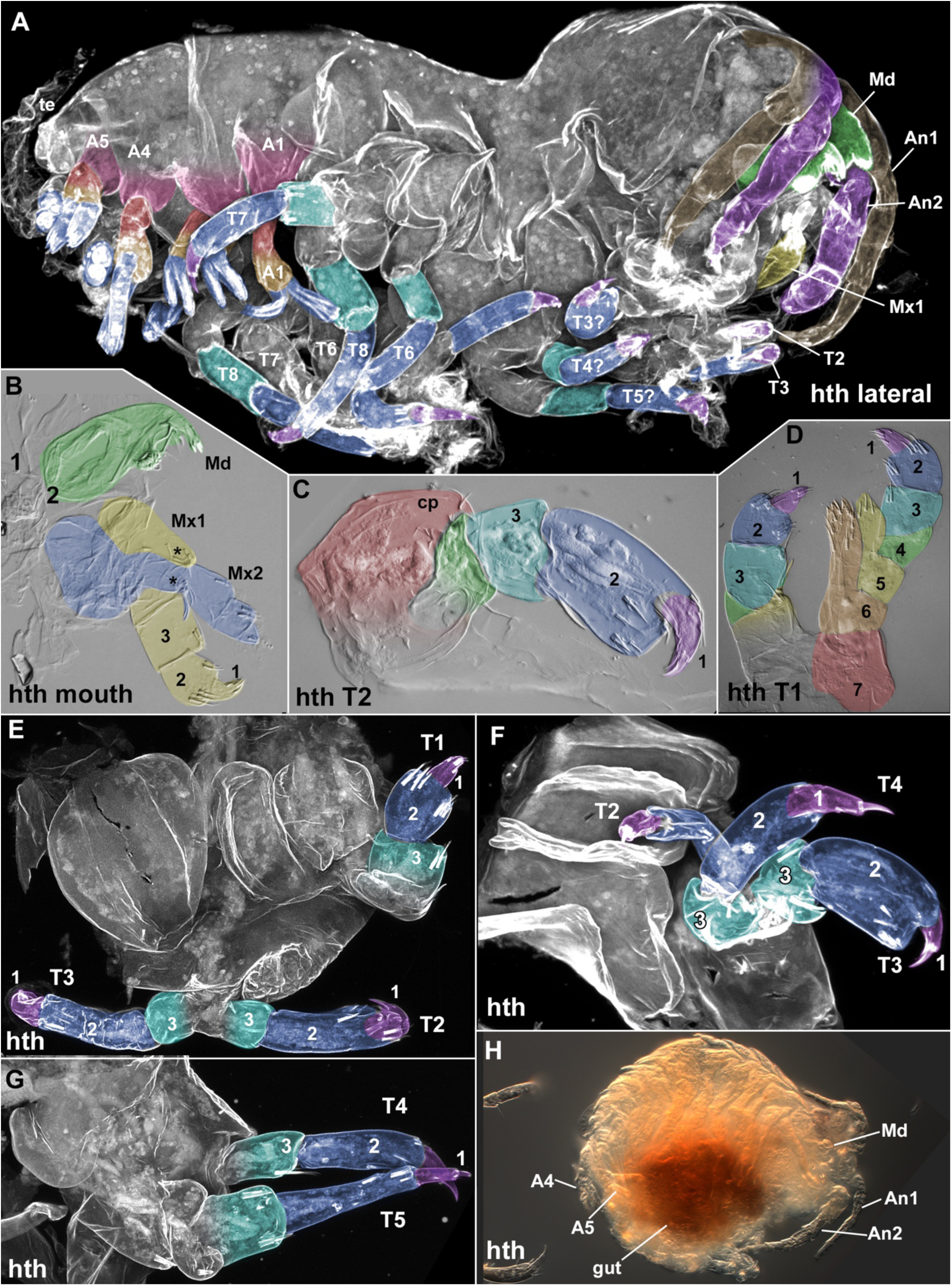
Homothorax phenotypes. Proximal leg segments deleted, leaving ls1, 2, and the distal part of ls3 normal. Colors and numbers in head as in Figure 1 legend, colors and numbers in thoracic and abdominal leg segments as in Fig. 2 legend. A, whole hatchling, oblique view. Legs labeled where identifiable. Mobile part of abdominal swimmerets and uropods appear normal, despite emerging from a lateral body wall (=ls 8 = precoxa) that is fused and distorted. B, dissected mouthparts, mandible (Md, green), maxilla 1 (Mx1, yellow), maxilla 2 (Mx2, blue). Mx1 and Mx2 transformed towards generic uniramous identity with loss of endites (*) and gain of joints and distal leg segments (ls1 and ls2 discernible). C, dissected T2, proximal leg segments deleted. D, dissected T1, proximal leg segments of left T1 deleted, right T1 appears wild type. E, dissected T1-3, proximal leg segments deleted, and base of T2 and T3 fused together. F, dissected T2-4, proximal leg segments deleted, and base of legs fused together. G, dissected T4, 5, proximal leg segments deleted, and base of legs fused together. H, gut defect, body segments reduced, leading to a short and round body. The gut appears shortened and red-brown.

**Figure 11.**
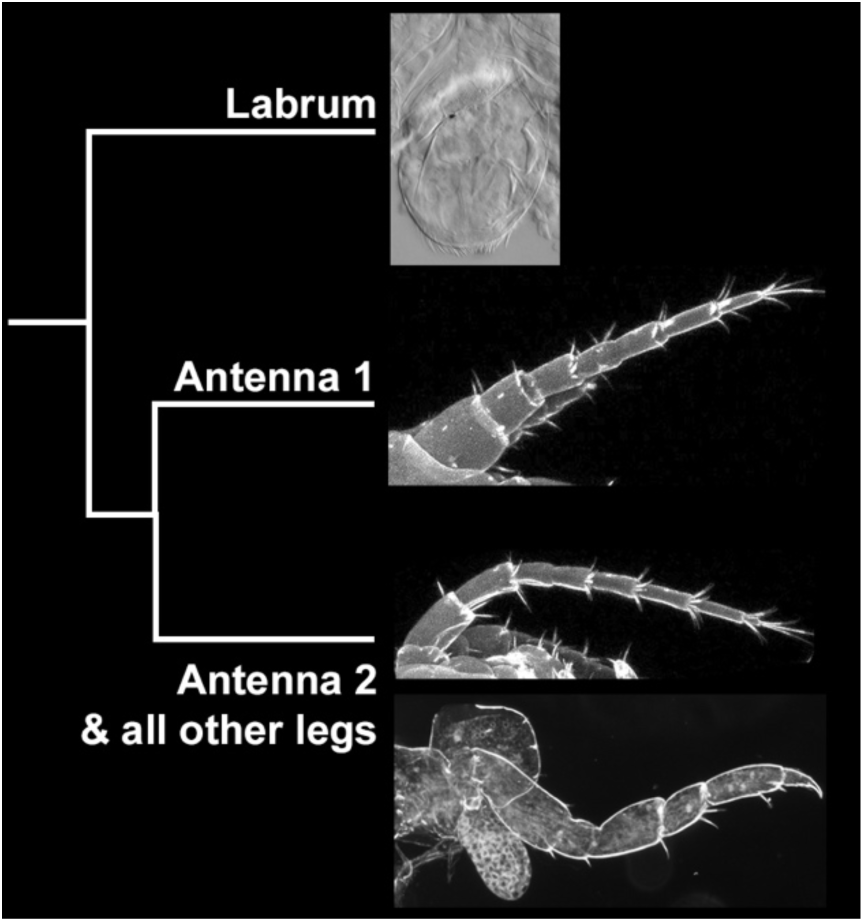
Labrum and canonical legs could all be derived from an ancient proto-limb that expressed the leg gene network. These proto-limbs subsequently diverged in function and morphology. The anterior labrum would represent the most ancient outgroup leg type with a more ancient wiring of the leg gene network and segmentation gene network. An1 and all other canonical legs subsequently diverged from the labrum, then An2 and all other canonical legs diverged from An1.

In the biramous abdominal appendages (A1 - 3 swimmerets and A4 - 6 uropods), the mobile base of the leg is considered to consist of a fused ls6 and ls7 (basis and coxa), and the phenotypes for *Sp6-9*, *Dll*, and *dac*2 agree with this interpretation. It is therefore curious that loss of *exd* or *hth* do not seem to affect this presumed proximal mobile base, but do affect the proximal-most leg segment, ls8 (=precoxa = lateral body wall = lateral tergum) that this mobile base stands on. This could possibly be explained by a lack of function of *exd* or *hth* in this mobile base despite their expression here, or some peculiarity of biramous appendages.

Interestingly, *exd* is expressed more strongly in the endopod than the exopod, which could perhaps be related to the difference in *Dll* phenotype between exopod and endopod (Fig. S3).

While the proximal mobile base of the biramous legs is difficult to interpret, the phenotypes for *Sp6-9* and *Dll* suggest that the 2-segmented exopod of A1 - 3 swimmerets is composed of a short ls1 and long 2 (*dac*tyl and propodus). Given that ls2 is the longer of the two exopod segments, the similar looking long segment on the endopod would likely correspond to ls2.

In milder *exd* or *hth* phenotypes, discrete regions of red pigment form on the antennae and T1 (Fig4.14). These regions may represent ectopic eyes. Ectopic eye formation is also seen when *Exd* and *Hth* are knocked out in *Drosophila* (González-Crespo & Morata, 1995; Inbal et al., 2001; Pai et al., 1998). Intriguingly, the pharynx also appears to form a large ectopic eye.

Rows of fine bristles identify this region as the gastric mill of the pharynx. There do not seem to be reports of eyes forming in the pharynx in *Drosophila*, but this could be due to the experimental mutation used, or understandably, because no one has looked in the *Drosophila* pharynx for ectopic eyes.

In conclusion, the function of leg patterning genes like *Dll*, *Sp6-9*, *dac*, *exd*, and *hth*, in conjunction with the expression of *ara*, provide useful readouts of arthropod appendage composition, especially in highly modified legs like the mouthparts. This approach could also be applied to deciphering the composition and homologies of other highly modified legs, like arthropod genitalia.

## METHODS

### MICROINJECTION AND RAISING

Embryos were injected in filtered artificial seawater (FASW) at either the 1-cell or 2-cell stage, the latter of which gives rise to roughly the left and right halves of the animal. Injected embryos were raised in Petri dishes at 26°C, and transferred to new FASW and a new Petri dish every day until hatching. After hatching (or dissection, if they were too sick to hatch), hatchlings were incubated at 4°C overnight to relax their muscles and discourage curling up during fixation. Hatchlings were then added to 4% PFA in FASW at 4°C, and fixed overnight at 4°C. The following day, hatchlings were washed 3 x 5 minutes in PBTween, and put into 50% then 70% glycerol in 1xPBS.

### HATCHLING DISSECTION AND IMAGING

Whole hatchlings in 70% glycerol were placed ventral-side down in a glass-bottom Petri dish and held in place with two pieces of tungsten wire for confocal imaging on a Zeiss 700.

Hatchlings are autofluorescent in the 405nm DAPI channel, so no dyes were needed. Hatchlings in 70% glycerol were dissected with sharpened tungsten needles in a 3-well glass dish.

Individual dissected legs were mounted on slides and imaged on a Zeiss Axio compound microscope in DIC and dark field.

### TRADITIONAL COLORIMETRIC IN SITU

Cloning and RNA probe synthesis. Total RNA was extracted from a large pool of Parhyale embryos at multiple stages of embryogenesis, from stages 12–26 using TRIzol. Complimentary DNA was generated using SuperScript III. Primers were generated with Primer3 (http://bioinfo.ut.ee/primer3-0.4.0), with a preferred product size of 700 base pairs, and we avoided evolutionarily conserved domains to avoid possible probe cross-reactivity. Inserts were amplified with Platinum Taq (Thermo Fisher Scientific; 10966026), ligated into pGEM-T Easy vectors (Promega; A1360) and transformed into Escherichia coli. The resulting plasmids were cleaned with a QIAprep Miniprep kit (Qiagen; A1360) and sequenced to verify the correct insert and determine sense and anti-sense orientation. In situ templates were generated by PCR from these plasmids using M13F/R primers and purified with a Qiagen PCR Purification kit (Qiagen; 28104). The resulting PCR products were used to generate digoxigenin-labelled RNA probes (Roche; 11175025910) using either T7 or Sp6 RNA polymerase. RNA probes were precipitated with LiCl, resuspended in water and run on an agarose bleach gel (Aranda et al., 2012) to check that the probes were the correct size, and the concentration was determined using a NanoDrop 1000. The probes were used at a concentration of 1–5 ng /μl. In situ protocol. Embryo collection, fixation and dissection were performed as previously described71. In situ hybridization was performed as previously described(Rehm et al., 2009). In brief, embryos were fixed in 4% paraformaldehyde in artificial seawater for 45 min, dehydrated to methanol and stored overnight at −20 °C to discourage the embryos from floating during the later hybridization solution step.

The embryos were rehydrated using 1x phosphate-buffered saline (PBS) with 0.1% Tween 20 (PBST), post-fixed for 30 min in 9:1 PBST:paraformaldehyde and washed in PBST. The embryos were incubated in hybridization solution at 55 °C for at least 36 h. The embryos were blocked with 5% normal goat serum and 1x Roche blocking reagent (Roche; 11096176001) in PBST for 30 min. Sheep anti-DIG-AP antibody (Roche; 11093274910) was added at 1:2,000 and incubated for 2 h at room temperature. The embryos were developed in BM Purple (Roche; 11442074001) for a few hours or overnight. After the embryos were sufficiently developed, they were dehydrated using methanol to remove any pink background, then rehydrated using PBST. The embryos were then moved to 1:1 PBS:glycerol with 0.1 mg ml−1 4′,6-diamidino-2- phenylindole, then 70% glycerol in PBS.

### IN SITU HCR V 3.0

In situ HCR version 3.0 from Molecular Instruments and Choi 2018(Choi et al., 2018).

Parhyale and Tribolium complimentary DNA sequences were submitted to Molecular Instruments and the probe sets are available from the company. In situ hybridizations were performed using their whole-mount Drosophila embryo protocol, with the following exceptions: (1) the embryos were left on the bench, not rocked back and forth; (2) the embryos were permeabilized in sodium dodecyl sulfate detergent solution (1% sodium dodecyl sulfate, 0.5% Tween 20, 50 mM Tris-HCL (pH 7.5), 1 mM EDTA (pH 8.0) and 150 mM NaCl) for 30 min instead of proteinase K, which improved the morphology in our hands. The probe set numbers from Molecular Instruments are provided in Bruce & Patel 2020 (Bruce & Patel, 2020).

### CRISPR–CAS9 GUIDE RNA GENERATION, INJECTION AND IMAGING

Guide RNAs were generated using ZiFit(Sander et al., 2007, 2010), as previously described(Martin et al., 2016). Single guide RNAs (sgRNAs) were ordered from Synthego. The injection mixes had a final concentration of 333 ng μl−1 Cas9 protein, 150 ng μl−1 sgRNA (for both single-and double-guide injection mixes) and 0.05% phenol red for visualization during injection, all suspended in water. One-or two-cell embryos were injected with approximately 40–60 pl sgRNA mixture, as previously described(Martin et al., 2016). The resulting knockout hatchlings were fixed in 4% paraformaldehyde in artificial seawater at 4 °C for 1–2 d, then moved to 70% glycerol in 1x phosphate buffered saline (PBS). Dissected hatchling limbs were visualized with Zeiss LSM 700 and 780 confocal microscopes using 405nm ultraviolet autofluorescence to visualize the cuticle. Z-stacks were assembled with Volocity version 6.0 (PerkinElmer). Adobe Photoshop CS6 or 2023 was used to create color-overlays on hatchlings using the Overlay command and HCR expression channels in embryos were combined using the Screen function.

## AUTHOR CONTRIBUTIONS

H.S.B. conceived, designed, and performed the experiments and wrote the manuscript. N.H.P. edited the manuscript.

---

**Figure S1.**
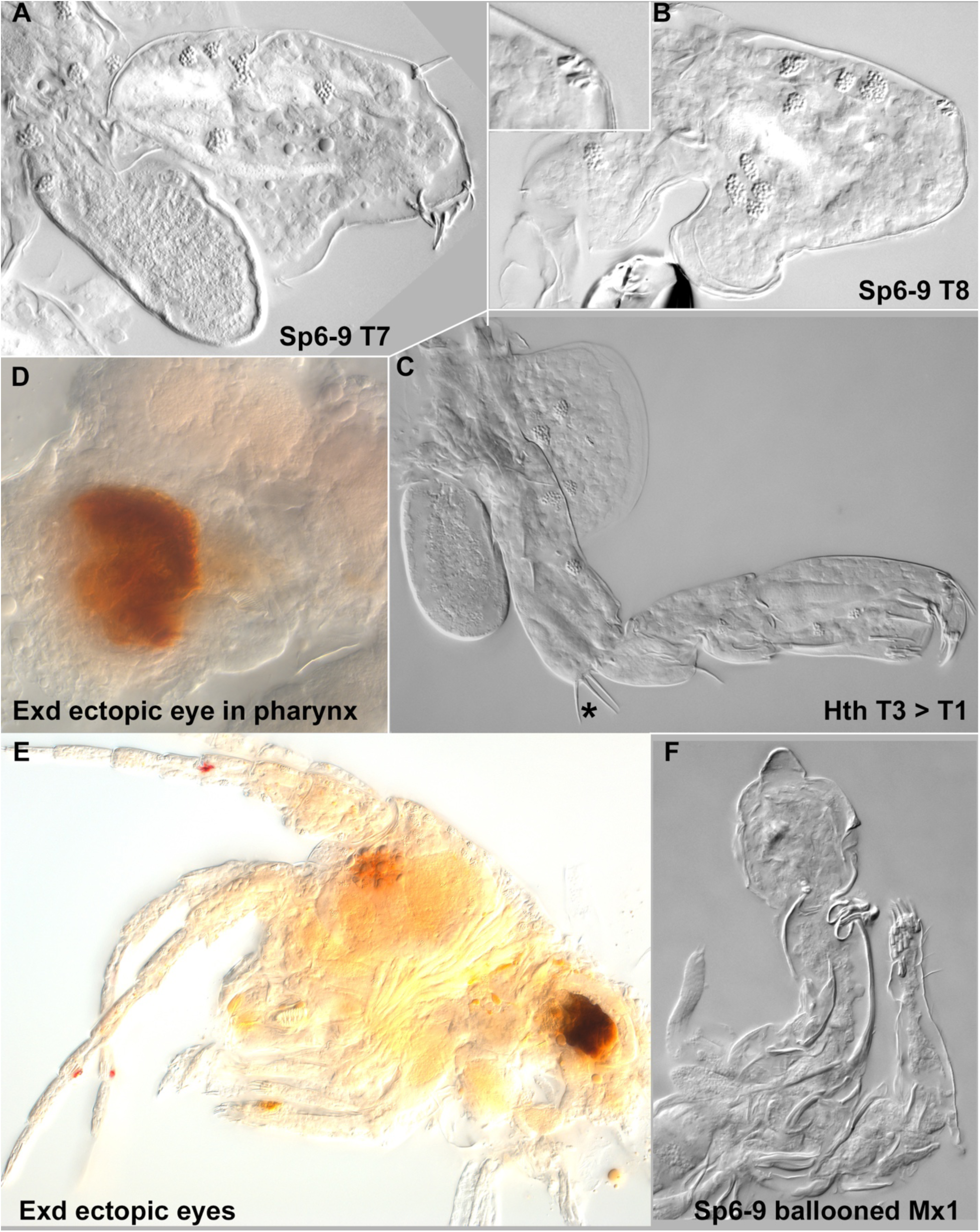
Additional phenotypes. A, *Sp6-9* knockout in T7 with possible ectopic exopod. B. *Sp6-9* knockout in T8 basal plate with narrow neck, possibly transformed towards coxal plate. C, *hth* knockout in T2 transformed towards T1 identity with endite (*). D, *exd* knockout with ectopic eye in pharynx. E, *exd* knockout with ectopic eyes. F, *Sp6-9* knockout in maxilla 1 causes ballooned tissue and deletes one of the endites in maxilla 2.

**Figure S2.**
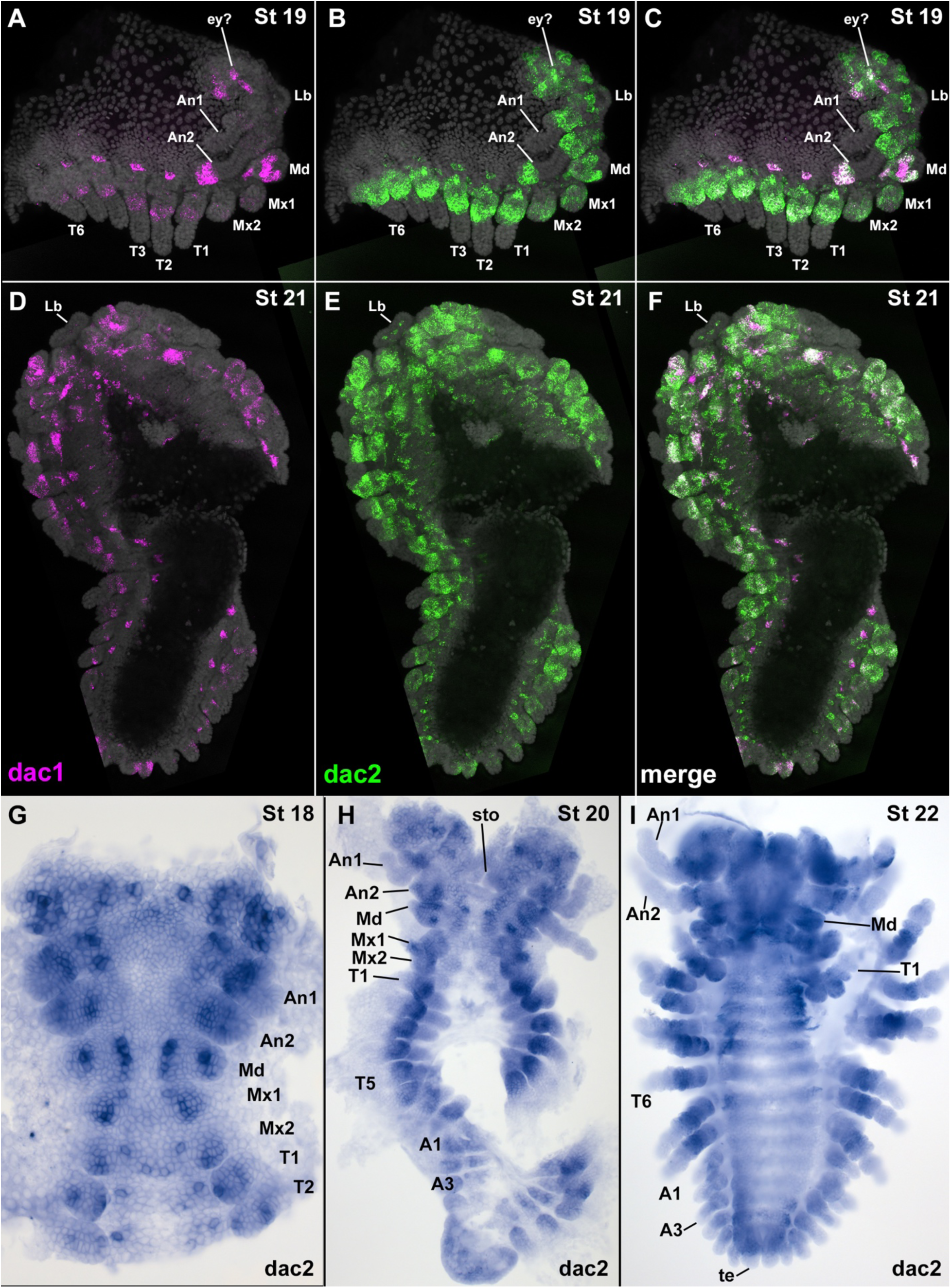
Expression of *dac*1 and *dac*2. A-F, *dac*1 and *dac*2 have similar but not identical expression domains in the medial leg and in a spot on the anterior side of each proximal leg. *dac*1is weaker in the medial leg and stronger in a proximal spot, while *dac*2 is stronger in the medial leg and weaker in the proximal spot. A, *dac*1 expression in Stage 19 embryo. B, *dac*2 expression in Stage 19 embryo. C, merge of *dac*1 and *dac*2 in Stage 19 embryo. D, *dac*1 expression in Stage 21 embryo. E, *dac*2 expression in Stage 21 embryo. F, merge of *dac*1 and *dac*2 in Stage 21 embryo. G, *dac*2 expression in Stage 18 embryo. H, *dac*2 expression in Stage 20 embryo. I, *dac*2 expression in Stage 22 embryo.

**Figure S3.**
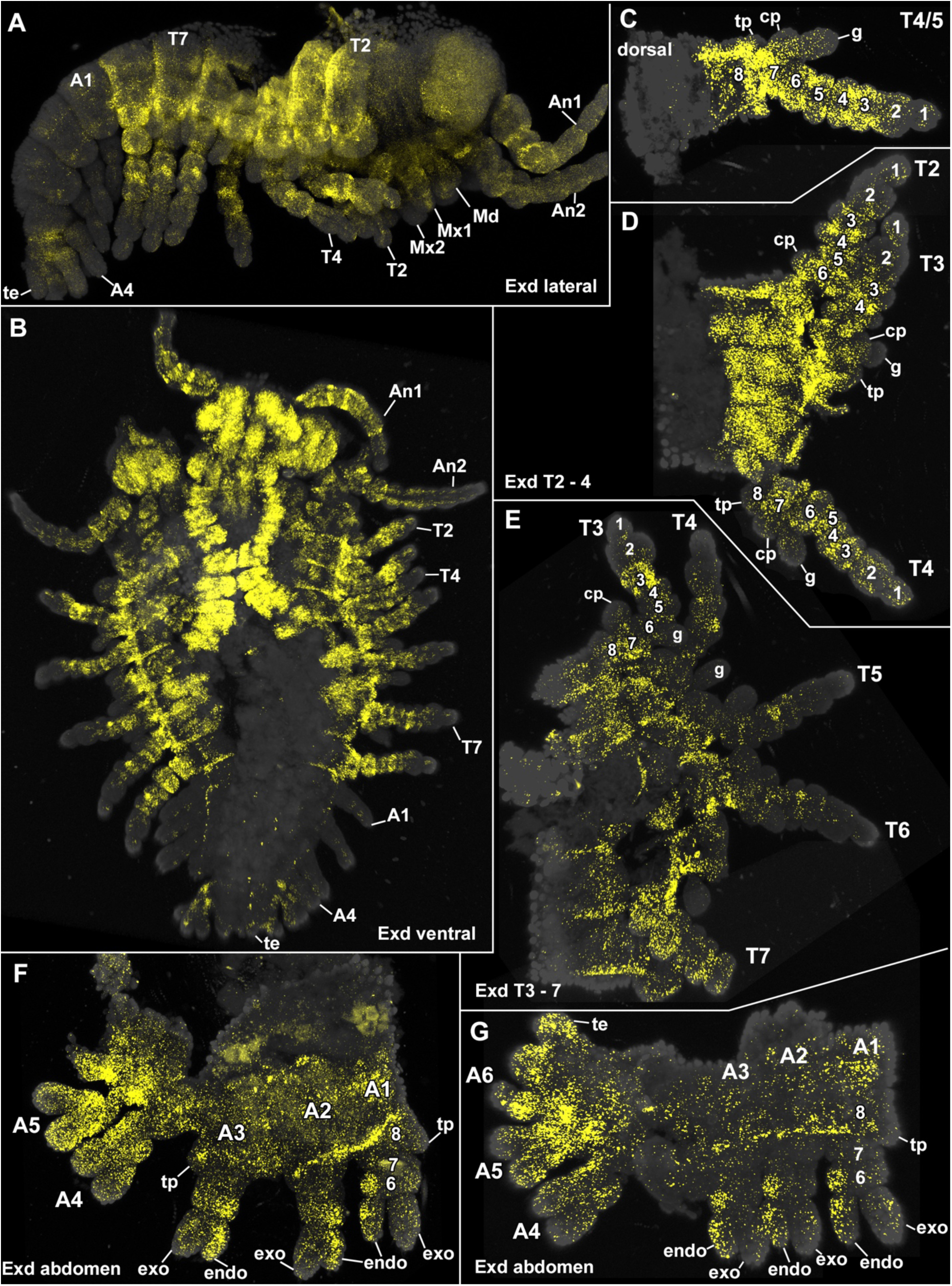
Expression of extradenticle. Extradenticle is expressed in the proximal leg, and what appears to be the leg-derived lateral body wall, but not in the medial, non-leg-derived body wall. Expression is weaker or absent in ls1 and ls2. Expression appears weaker in the abdomen, especially in the A1-3 swimmerets. A, whole embryo, lateral view. B, whole embryo, ventral view. C. Dissected T4 or T5 leg. D, dissected T2-4 legs and non-leg-derived body wall. E, dissected T3-7 legs and non-leg-derived body wall. F, dissected abdomen A1-5. *Exd* is stronger in the endopod than exopod in the A1-3 swimmerets. G, dissected abdomen A1-6 and telson (te). *Exd* is stronger in the endopod than exopod in the A1-3 swimmerets.

**Figure S4.**
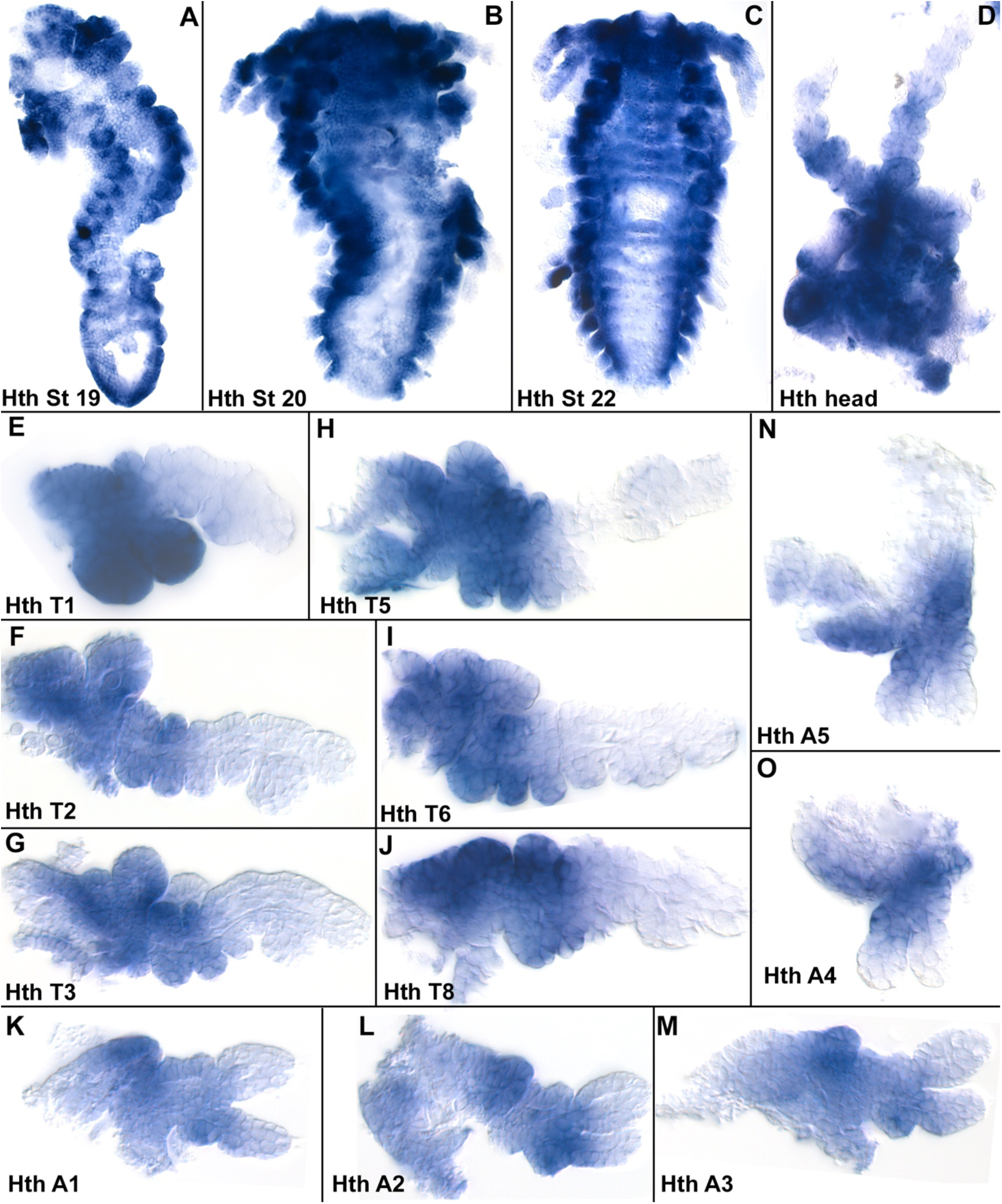
Expression of homothorax. A, Stage 19. *Hth* is not expressed in the mandible. B. Stage 20. *Hth* is expressed in the proximal legs, but not in the telson. C, Stage 22. *Hth* is expressed in the lateral edges of the labrum. D, dissected head. *hth* is expressed in the lateral edges of the labrum, and proximal An1 and An2. E, T1. *Hth* is expressed in proximal leg segments through ls5 including the endites on ls5 and ls6. F, T2. *Hth* is expressed in proximal leg segments through ls5, but not in the gill. G, T3. H, T5. I, T6. J, T8. K, A1. L, A2. M, A3. N, A5. O, A4.

**Figure S5.**
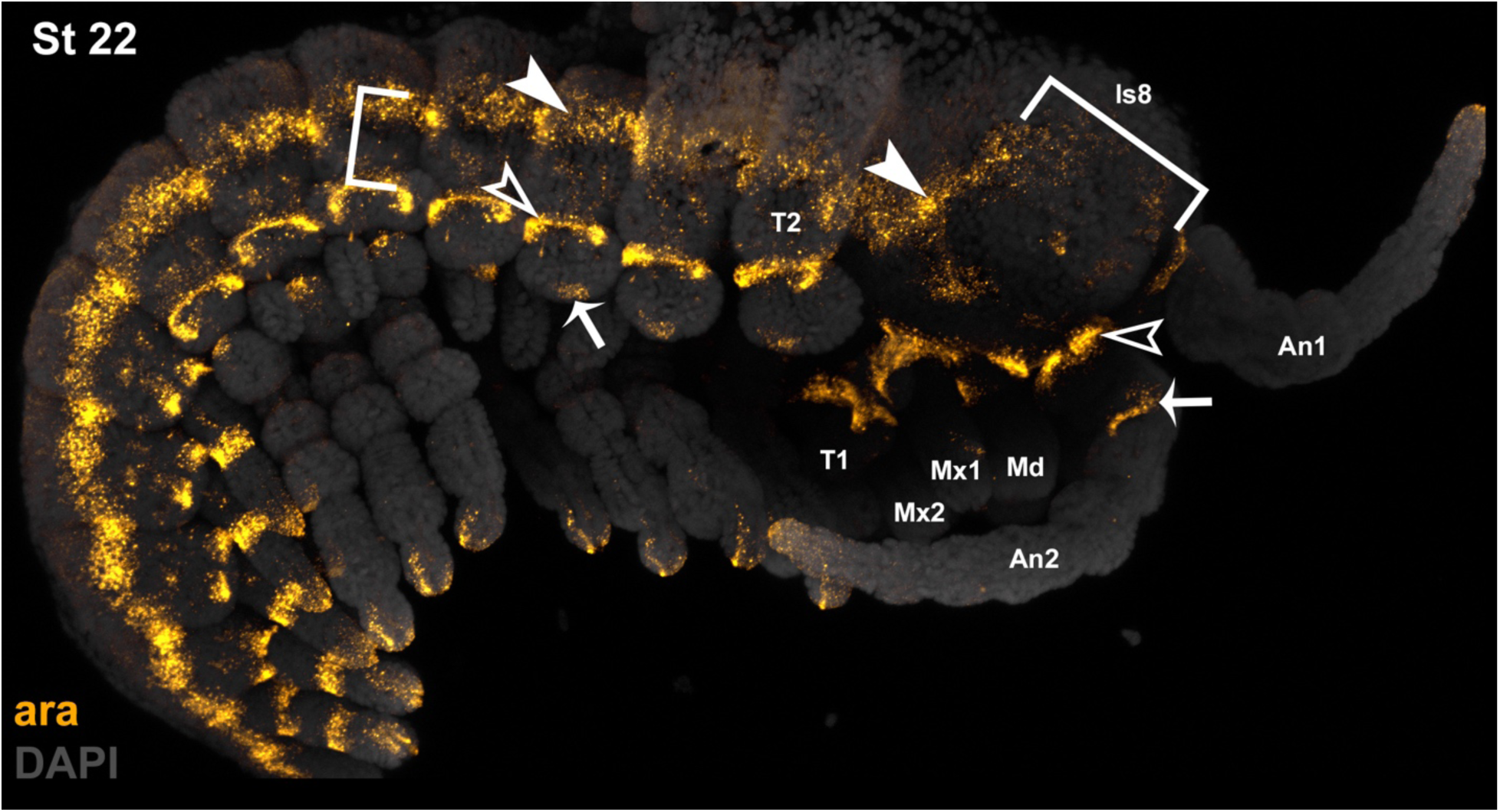
Expression of *araucan* in the *Parhyale* head suggests that most of lateral head mass is a conglomerate of multiple ls8’s (precoxae) from the head appendages. The precoxae presumably surround the base of each coxa circumferentially, both laterally and medially to each coxa. Given that the paragnaths resemble and function similar to endites, and emerge from the mandibular “sternum”(Wolff & Scholtz, 2006), here proposed to be the precoxa (ls8), the paragnaths may represent to the endites of the Md precoxa.

